# Scale cortisol levels vary with habitat type and population in the Mexican tetra *Astyanax mexicanus*

**DOI:** 10.1101/2025.03.09.642216

**Authors:** Marc B. Bauhus, Luise Wander, David Boy, Ana Santacruz, Ernesto Maldonado, Joachim Kurtz, Sylvia Kaiser, Robert Peuß

**Affiliations:** Institute for Evolution and Biodiversity, University of Münster, Germany; Institute for Integrative Cell Biology and Physiology, University of Münster, Germany; Stowers Institute for Medical Research, Kansas City, MO, USA; EvoDevo Research Group, Unidad Académica de Sistemas Arrecifales, Instituto de Ciencias del Mar y Limnología, Universidad Nacional Autónoma de México, Puerto Morelos, Quintana Roo, Mexico; Joint Institute for Individualisation in a Changing Environment, University of Münster and Bielefeld University, Münster, Bielefeld, Germany; Department of Behavioural Biology, University of Münster, Germany

**Author notes:** Corresponding author: Robert Peuß, Institute for Integrative Cell Biology and Physiology, Schlossplatz 8, 48143 Münster, Germany, Mail.

**Keywords:** *Astyanax mexicanus*, environmental variability, scale cortisol, stress

## Abstract

Cortisol is a key hormone involved in stress responses, and its regulation reflects how organisms cope with environmental stressors. While acute stress responses are well studied, much less is known about how cumulative, long-term stress shapes the responses of populations adapted to different habitats. Here, we investigated scale cortisol – a biomarker of long-term stress exposure in fish – in wild and laboratory reared populations of *Astyanax mexicanus*, a species with surface and cave-dwelling ecotypes adapted to contrasting environments. We measured scale cortisol levels in six populations (four cave and two surface) over two consecutive years, alongside abiotic parameters in the respective habitats. Some cave populations exhibited lower scale cortisol levels than surface fish under both wild and laboratory conditions, suggesting a genetically based enhancement of stress-coping ability or altered cortisol-mediated metabolic and osmoregulatory processes. Moreover, populations experiencing more environmental variability – such as the lake surface habitat and hydrologically dynamic cave systems – showed higher scale cortisol levels and increased inter-individual variation. These results suggest that evolutionary adaptations to different habitats shape long-term stress physiology in *A. mexicanus*, and they highlight scale cortisol as a valuable tool for assessing how organisms physiologically cope with diverse and extreme environments.

## Introduction

Stress is a response to environmental challenges or threats (stressors), triggering a complex physiological and behavioral reaction that enables individuals to cope with environmental changes (Schulte et al., 2014). Stress can be induced by changes in abiotic conditions such as temperature (Gates et al., 1992), oxygen levels (Van Raaij et al., 1996) and pH (Wang et al., 2009) or by biotic factors such as predators (Van Dievel et al., 2016), parasites (Huffman et al., 2004), aggressive interactions among conspecifics (Sachser and Lick, 1991), and food availability (Pravosudov et al., 2001). During the vertebrate stress response, the adrenal glands (or interrenal tissue in fish) release glucocorticoids, for example cortisol or corticosterone, which maintain homeostasis by regulating, among other things, metabolism (Brillon et al., 1995; Christiansen et al., 2007; Vijayan et al., 1997) and immunity (Webster Marketon and Glaser, 2008).

Since habitats differ widely in abiotic and biotic characteristics, glucocorticoid regulation can vary across individuals and populations, reflecting both genetic adaptations and plastic responses to environmental pressures (DeKoning et al., 2004; Schulte et al., 2014). However, whether contrasting habitat types – such as different surface *vs.* cave water bodies – exert long-term and cumulative effects on an organism’s stress physiology remains unclear. To investigate this, we use the Mexican Tetra *Astyanax mexicanus* which depicts an extreme example of local adaptation to different habitats. *A. mexicanus* consists of two ecotypes: surface-dwelling (surface fish) and cave-dwelling (cavefish), both distributed in various caves and surface water bodies across Northeastern Mexico (Mitchell et al., 1977) (Fig. 1A). Cave aquatic environments present stressors such as reduced and infrequent availability of nutritional resources due to the absence of primary energy production in total darkness (Poulson, 2001; Venarsky et al., 2014). However, they also provide more stable abiotic conditions (Krishnan et al., 2020; Poulson and White, 1969; Tabin et al., 2018) but with differences between caves and pools within caves depending on the hydrological regime (Legendre et al., 2026). Consequently, cavefish have evolved several adaptations, some of which are population dependent, that allow them to thrive in resource-limited conditions. These include metabolic adaptations such as elevated blood sugar levels and insulin resistance (Riddle et al., 2018; Shimobayashi et al., 2023), enhanced lipogenesis (Xiong et al., 2022) and increased fat accumulation (Xiong et al., 2018). Behavioral adaptations include binge eating (Aspiras et al., 2015), reduced sleep (Yoshizawa et al., 2015) and decreased aggression (Elipot et al., 2013; Rodriguez-Morales et al., 2022). Additionally, cavefish must cope with hypoxia (Krishnan et al., 2020; Ornelas-García et al., 2018; Rohner et al., 2013) which has led to an increase in embryonic erythrocyte production (van der Weele and Jeffery, 2022) as well as larger erythrocytes with higher hemoglobin content per cell in adult fish (Boggs et al., 2022).

**Figure 1.**
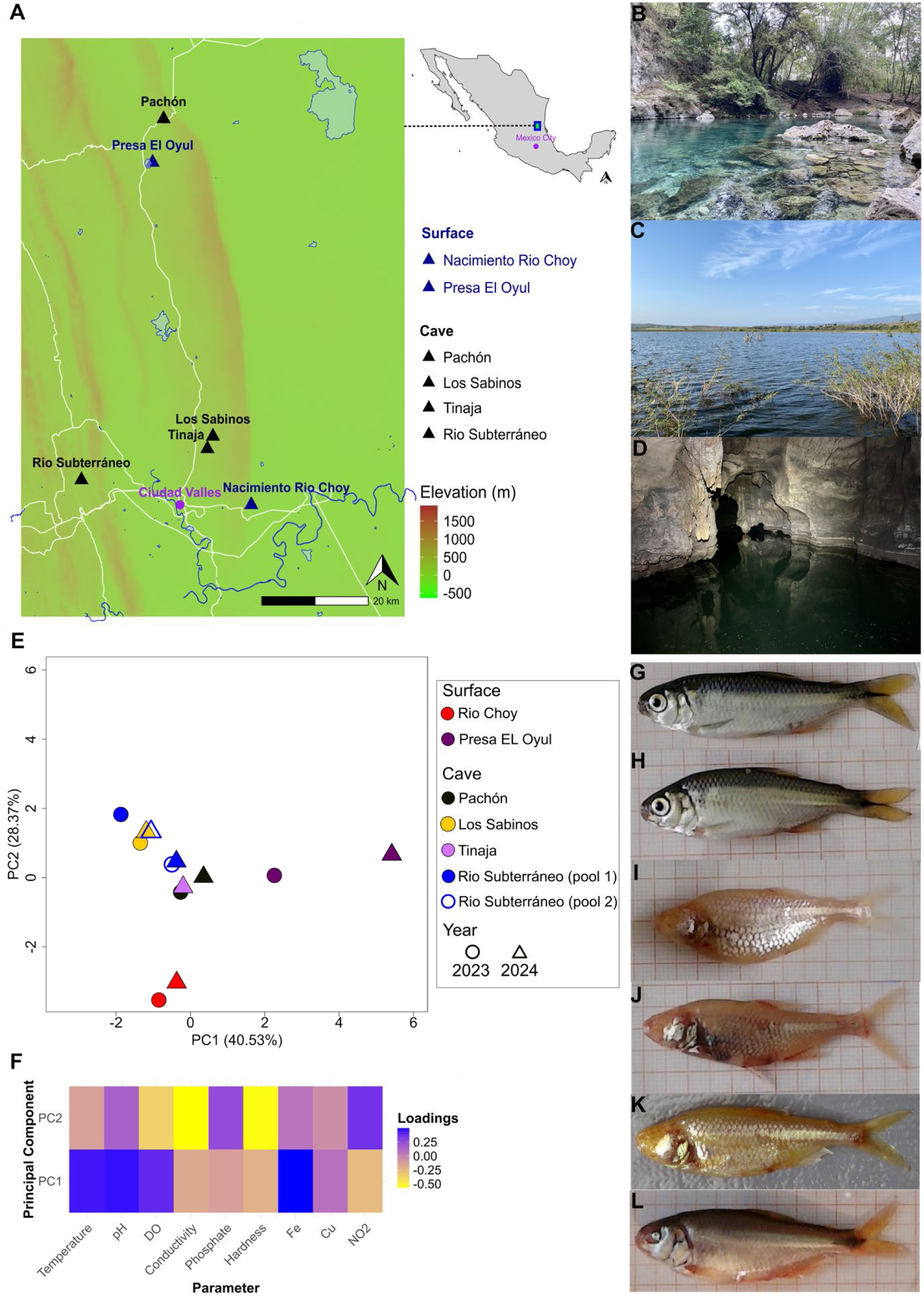
Sampling habitats. (A) Map of sampling area in the Sierra de El Abra Tanchipa and Sierra Micos region in North-eastern Mexico with sampling locations of respective populations. (B) Stream surface fish habitat (Nacimiento Rio Choy). (C) Lake surface fish habitat (Presa El Oyul). (D) Cavefish habitat (Pachón) cave. (E) Principal component analysis (PCA) plot based on abiotic parameters from Table 1 showing scores of principal components (PC) 1 (x-axis) and 2 (y-axis) per habitat and year. The percentage of explained variation per PC is displayed in brackets. (F) Heatmap showing PC loading factors per measured parameter. Representative images of (G) Rio Choy and (H) Presa El Oyul surface- and (I) Pachón, (J) Los Sabinos, (K) Tinaja and (L) Rio Subterráneo cavefish.

**Table 1.** Abiotic water parameters in surface (Rio Choy, Presa El Oyul) and cave (Pachón, Los Sabinos, Rio Subterráneo, Tinaja) habitats measured in spring 2023 and 2024.

| Site | Date | Temperature [C°] | pH | Dissolved Oxygen [mg/L] | Conductivity [μs/cm] | Phosphate [mg/L] | Hardness (CaCO<sub>3</sub>) [mg/L] | Iron [mg/L] | Copper [mg/L] | Nitrite [mg/L] |
| --- | --- | --- | --- | --- | --- | --- | --- | --- | --- | --- |
| Rio Choy | 09.04.2023 | 25.9 | 7.0 | 7.36 | 1164 | 0.75 | 710.5 | 0 | 0 | 0.004 |
| Rio Choy | 11.03.2024 | 25.8 | 7.2 | 6.80 | 1123 | 0.06 | 560 | 0 | 0.02 | 0.005 |
| Presa El Oyul | 14.04.2023 | 26.0 | 8.4 | 7.59 | 428 | 1.00 | 214 | 0.05 | 0.04 | 0.002 |
| Presa El Oyul | 17.03.2024 | 30.5 | 9.5 | 9.40 | 316 | 0.60 | 64 | 0.13 | 0 | 1E-10 |
| Pachón | 06.04.2023 | 24.3 | 7.3 | 5.34 | 568 | 0.45 | 307 | 0.005 | 0.005 | 0.013 |
| Pachón | 17.03.2024 | 25.7 | 7.5 | 5.80 | 490 | 0.60 | 208 | 0 | 0.01 | 0.027 |
| Los Sabinos (pool 1) | 12.04.2023 | 22.8 | 7.4 | 1.97 | 484 | 1.30 | 251 | 0 | 0 | 0.065 |
| Los Sabinos (pool 1) | 21.03.2024 | 23.1 | 7.8 | 1.73 | 534 | 0.60 | 239 | 0 | 0.03 | 0.177 |
| Tinaja (pool 2) | 19.03.2024 | 23.3 | 8.1 | 6.29 | 667 | 2.40 | 346 | 0 | 0 | 0.009 |
| Rio Subterráneo (pool 1) | 07.04.2023 | 23.7 | 7.2 | 1.76 | 470 | 1.30 | 232 | 0 | 0 | 0.229 |
| Rio Subterráneo (pool 2) | 07.04.2023 | 23.8 | 7.3 | 5.77 | 481 | 1.40 | 228 | 0 | 0 | 0.048 |
| Rio Subterráneo (pool 1) | 14.03.2024 | 25.0 | 7.5 | 3.37 | 524 | 1.90 | 234 | 0 | 0 | 0.007 |
| Rio Subterráneo (pool 2) | 14.03.2024 | 25.1 | 7.2 | 3.38 | 519 | 4.70 | 212 | 0 | 0 | 0.063 |

Unlike cavefish, surface-dwelling *A. mexicanus* experience greater fluctuations in abiotic conditions, such as temperature variation (Tabin et al., 2018), and are exposed to more biotic stressors, including parasites (Peuß et al., 2020; Santacruz et al., 2023, 2020), predators (Chin et al., 2018; Mitchell et al., 1977; Yoshizawa and Library, 2015) and aggressive interactions among conspecifics (Elipot et al., 2013; Rodriguez-Morales et al., 2022).

Strong ecological differences between cave and surface habitats resulted in specific stress response adaptations of cave- and surface fish. For example, cavefish show lower plasma cortisol levels upon exposure to confined space (Gallo and Jeffery, 2012), lower whole-body cortisol in total darkness (Bilandžija et al., 2020), and reduced stress behaviors such as freezing, and prolonged bottom dwelling (Chin et al., 2018; Padmanaban et al., 2025) compared to surface fish.

In addition to their ecological divergence, cavefish differ from surface fish in key life-history traits–including slower growth rates (Simon et al., 2017), increased offspring investment under nutrient limitation (Xia et al., 2025), and year-round reproduction (Espinasa et al., 2023)–, traits known to interact with glucocorticoid release (Crespi et al., 2013).

While short-term elevations in blood-plasma cortisol occur in response to acute stress, scale cortisol levels provide a measure of long-term experienced stress, as cortisol accumulates in scales over time (Aerts et al., 2015). Consequently, scale cortisol levels have been widely used to assess long-term experienced stress across various fish species (Britton et al., 2023; Lebigre et al., 2022; Roque d’orbcastel et al., 2021; Vercauteren et al., 2022).

To examine differences in long-term experienced stress between surface and cave-adapted *A. mexicanus*, we measured cortisol levels in scales of wild surface and cave individuals from different populations across two consecutive years. We sampled two distinct surface populations: Rio Choy (a river population; Fig. 1B, G) and Presa El Oyul (a lake population; Fig. 1C, H) – as well as four cave populations: Pachón (Fig. 1D, I), Los Sabinos (Fig. 1J), Tinaja (Fig. 1K) and Rio Subterráneo (Fig. 1L) – which represent an isolation gradient from surface habitats (Mitchell et al., 1977; Moran et al., 2023; Simon et al., 2017). Additionally, we measured several abiotic parameters in both years to characterize the abiotic environment of each habitat type. To disentangle environmental and genetic effects on cortisol levels, we compared scale cortisol levels from wild individuals of Rio Choy surface and Pachón cavefish to those of laboratory-reared individuals from the same populations. We hypothesized that scale cortisol levels–reflecting long-term experienced stress–and individual variation therein would be lower in cavefish compared to surface fish, due to reduced stressors or stress responsiveness in cave-adapted populations. Furthermore, we expected this difference to persist under laboratory conditions, suggesting a genetic basis for stress-related adaptations to the cave environment. Our alternative prediction was that there are no differences in scale cortisol levels between laboratory-bred cave- and surface fish due to a plastic stress regulation.

## Material & methods

### Field sample collection

We collected wild individuals (males and females) of *A. mexicanus* in 2023 and 2024 under permit no. SPARN/DGVS/03371/23 granted by the Secretaría de Medio Ambiente y Recursos Naturales (SEMARNAT) to Ernesto Maldonado. In 2023, we sampled fish from Rio Subterráneo (Micos), Pachón and Los Sabinos caves as well as from Nacimiento Rio Choy (stream) and Presa El Oyul (lake) between April 5^th^ and April 15^th^. Additionally, we collected fish from Tinaja cave between December 2^nd^ and 3^rd^. In 2024, we resampled Rio Subterráneo, Tinaja, Rio Choy and Presa El Oyul (only females were found) between March 9^th^ and March 19^th^.

The Pachón cave is the most isolated cave among the sampled caves due to its elevated cave entrance (210 m above sea level) (Mitchell et al., 1977) and hence has the most stable abiotic conditions, whereas the other caves are still regularly flooded during the rainy season (Legendre et al., 2026). In Rio Subterráneo, there is a regular introduction of surface fish due to a karst stream flooding into the cave during the rainy season which is thought to result in hybridization with the resident cavefish (Garduño-Sánchez et al., 2023; Moran et al., 2023). Consequently, Rio Subterráneo cavefish show variable and intermediate phenotypes such as reduced but not completely lost eyes and weakened pigmentation (see Fig. 1L).

For practical reasons, and to minimize impact on cave populations, capture techniques had to be adjusted to the local conditions. In cave environments, we used dip nets to capture fish, while in stream and lake habitats, we employed improvised fishing poles equipped with size 16 hooks (Gamakatsu, Japan) baited with tortilla or dried mango to catch surface fish. After capture, fish were kept alive until dissection and transported in plastic bags with oxygen supply by air bubblers to an improvised field lab. Here, we euthanized all fish with an overdose of MS-222 (0.5 g/L) and dissected them as quickly as possible, no longer than 24h post-capture (Supplemental Tab. 1). We weighed all fish to the nearest mg and took standardized pictures to determine the standard length of each fish to the nearest mm with ImageJ. Using a metal spatula, we carefully collected nearly all scales per individual and transferred them to pre-weighed 2 mL Eppendorf-tubes, which we immediately flash froze in liquid nitrogen. We stored the samples at -20 °C until further processing. The sex of each fish was determined by opening the body cavity and screening the reproductive organs.

Furthermore, we measured temperature [C°], pH (in 2023), dissolved oxygen [mg/L], conductivity [µS/cm], phosphate high range [mg/L], hardness (CaCO_3_) high range [mg/L], iron [mg/L], copper [mg/L], and nitrite [mg/L] with a Hach SL 1000 multiparametric device and the respective Chemkeys and probes. Measurements were taken once per habitat and year (except Tinaja in 2023, see exact dates in Tab. 1). For pH measurements in 2024, we used an Akozon Smart Sensor Pen Type PH Meter Portable Water Quality Tester Acidimeter PH818.

### Laboratory fish husbandry and sample collection

We obtained parental Pachón cavefish from the laboratory of Sylvie Retaux (Paris-Saclay Institute of Neuroscience) and Rio Choy surface fish from the laboratory of Horst Wilkens (University of Hamburg). These lines have been kept and bred in the lab for at least two decades (Pachón) (Bilandžija et al., 2020) or longer (Rio Choy) (Wilkens, pers. comm.). We grouped at least four males and two females together for breeding, following the methods described by Baumann and Ingalls (2022). We maintained offspring from both populations at 22°C and fed them daily with freshly hatched *Artemia* larvae until they reached approximately 10 mm in size. Between 10 and 30 mm, we provided food five times per week using Gemma Micro (Skretting, Norway). As they grew between 30 and 45 mm, we adjusted the feeding schedule to four times per week with Gemma Silk (Skretting, Norway) and once per week with frozen bloodworms. After reaching 45 mm, we fed them twice per week with Gemma Silk and once per week with frozen bloodworms. We housed adult fish (> 45 mm) in groups of at least six individuals in 14 L aquaria with recirculating tap water, temperature control, and mechanical, UV, and biological filtration (Vewa Tech, Hamm, Germany). We kept the fish at a photoperiod of 14/10 h light/dark and used Osram Lumilux Cool Daylight HO 39W/865 T5 lamps for tank illumination. We regularly monitored the water quality in the fish tanks by measuring pH, conductivity, nitrite, ammonium and phosphate (See Supplemental Tab. 2). To minimize aggression in surface fish (Baumann and Ingalls, 2022), we enriched Rio Choy surface fish tanks with hexagonal plastic structures. Since effects of environmental enrichment on cortisol release are related to reduced aggression and anxiety (Näslund et al., 2013; Zhang et al., 2020), enrichment was not needed for cavefish which lost these behaviors (Chin et al., 2018; Elipot et al., 2013; Padmanaban et al., 2025; Rodriguez-Morales et al., 2022). Due to slower growth rates of cavefish (Simon et al., 2017) and in order to obtain enough scale material from fish of approximately the same size, we euthanized fish at 10 months (Rio Choy) and 14 months (Pachón) of age using an overdose of MS-222 (0.5 g/L) in accordance with the permit T24.016UMS granted by the University of Münster. Replicating the field pipeline, we collected nearly all scales per individual with a metal spatula and transferred them to pre-weighed 2 mL Eppendorf tubes, storing them at -20°C until analysis.

### Extraction of cortisol from scales and determination of cortisol levels

Before processing, we thawed the scales and washed them three times by adding 1.2 mL of isopropanol to each tube, shaking them at 20 Hz for 3 minutes in a ball mill (MM 400, Retsch GmbH, Germany). We discarded the supernatant and allowed the scales to dry overnight in a fume cupboard before recording their dry weights. Next, we placed two 5 mm steel balls into each tube and pulverized the scales for 3 minutes at 30 Hz in the ball mill. We weighed up to 13 mg of powdered scales and added 1.3 mL of 100% methanol, maintaining a final concentration of 10 mg powdered scales/mL methanol. We incubated the samples overnight in methanol on an overhead shaker. The next day, we centrifuged the samples at 1,000 × *g* for 1 minute, transferred the supernatant to a new tube, and centrifuged it again at 20,000 × *g* for 10 minutes. We transferred 1 mL of the supernatant to a new vial and evaporated it in a vacuum concentrator. We then resuspended the cortisol residue in 250 µL of ELISA buffer provided by the assay kit (Neogen, product no. #402710, USA). We selected this assay kit based on previous validations in other fish species (Britton et al., 2023; Carbajal et al., 2018; Ding et al., 2024) and because it showed the best performance during our own validation trials with an average recovery rate of approximately 90% (for details about the assay validation see Supplemental Methods). We stored the samples at −20°C until performing the ELISA. Before measurement, we thawed and vortexed the samples. We determined cortisol concentrations in duplicate following the manufacturer’s instructions. To minimize potential batch effects, we randomly loaded each ELISA plate with samples from different populations.

### Statistical analysis

All statistical analyses were conducted in R (version 4.3.2). First, we performed a principal component analysis (PCA) on abiotic parameters that were measured during the respective sampling day to characterize the abiotic environment of cave and surface habitats. Zero values were imputed to 1E-10.

For statistical analysis of scale cortisol levels, after excluding some individuals (Rio Choy 2023=4, Rio Choy 2024=2, Presa El Oyul 2023=2, Presa El Oyul 2024=1, Pachón 2023=13, Pachón lab=1, Los Sabinos 2023=2, Tinaja 2023=15, Tinaja 2024=1, Rio Subterráneo 2023=4, Rio Subterráneo 2024=4) due to non-sufficient scale material and a technical problem in the laboratory, we obtained the following sample sizes per population: Rio Choy 2023: N=21, Rio Choy 2024: N=23, Rio Choy lab: N=10, Presa El Oyul 2023: N=23, Presa El Oyul 2024: N=24, Pachón 2023: N=7, Pachon lab: N=9, Los Sabinos 2023: N=23, Tinaja 2023: N=8, Tinaja 2024: N=10, Rio Subterráneo 2023: N=21, Rio Subterráneo 2024: N=16. We first assessed normality using Shapiro-Wilk tests and visualized distributions with histograms and density plots. Since cortisol levels were not normally distributed but strictly positive, and right-skewed, we fitted generalized linear models (GLMs) with a Gamma distribution and log-link function. Model selection was based on Akaike Information Criterion (AIC) values, model diagnostics (residual plots from the car package) (see Supplemental Fig. 1-3), and the biological meaningfulness of fixed effects. We analyzed habitat and population effects in separate models because populations are fully nested within habitats, and not all populations were sampled in both years. Using different models allowed us to test for an overall habitat difference and assess within and among population × year variation. Model 1 tested for an overall habitat effect on cortisol levels including habitat (river, lake and cave), Fulton’s *k*-index as a measure for body condition (Barton et al., 2002), which controls for the length of an individual, and sex as fixed effects. Model 2 tested for population differences in scale cortisol levels including population × year together with Fulton’s *k*-index and sex as fixed effects. With Model 3, we tested for interaction effects of origin (wild *vs.* laboratory) × population for Rio Choy surface and Pachón cavefish, again including *k*-index and sex as fixed effects. Model residuals were checked for normality and homoscedasticity. We conducted pairwise comparisons of estimated marginal means (EMMs) using the emmeans package and applied compact letter display (CLD) to summarize significant differences among habitats and populations. Additionally, we used bootstrap resampling (1,000 iterations) to compare individual cortisol variability across populations and years. All statistical tests were two-tailed, and significance was set at *p* < 0.05.

## Results

### Abiotic parameters

A principal component analysis (PCA) based on abiotic parameters (Tab. 1) revealed notable differences not only between cave and surface habitats but also between the river (Rio Choy) and lake (Presa El Oyul) surface habitats (Fig. 1E). Additionally, Presa El Oyul exhibited significant differences in some parameters between 2023 and 2024, primarily reflected in principal component (PC) 1 (year 2023: 2.257, year 2024: 5.416) (Fig. 1E). PCA loading factors indicated that the variation in PC 1 was primarily driven by temperature (0.455), pH (0.470), dissolved oxygen (0.396), and iron (0.498) while variation in PC2 was mostly determined by conductivity (-0.556) and hardness (-0.565) Fig. 1F).

### Habitat and population differences in scale cortisol levels

We fitted two generalized linear models (GLMs) with scale cortisol level as response, habitat (model 1) or population × year (model 2) as the predictor and body condition (Fulton’s K-index) and sex as covariates. We found significantly higher scale cortisol levels in lake surface fish compared to river surface fish and cavefish while there was no difference between river surface and cavefish (Fig. 2A, see Supplemental Tab. 3, 4 for EMMs and pairwise comparisons).

**Figure 2.**
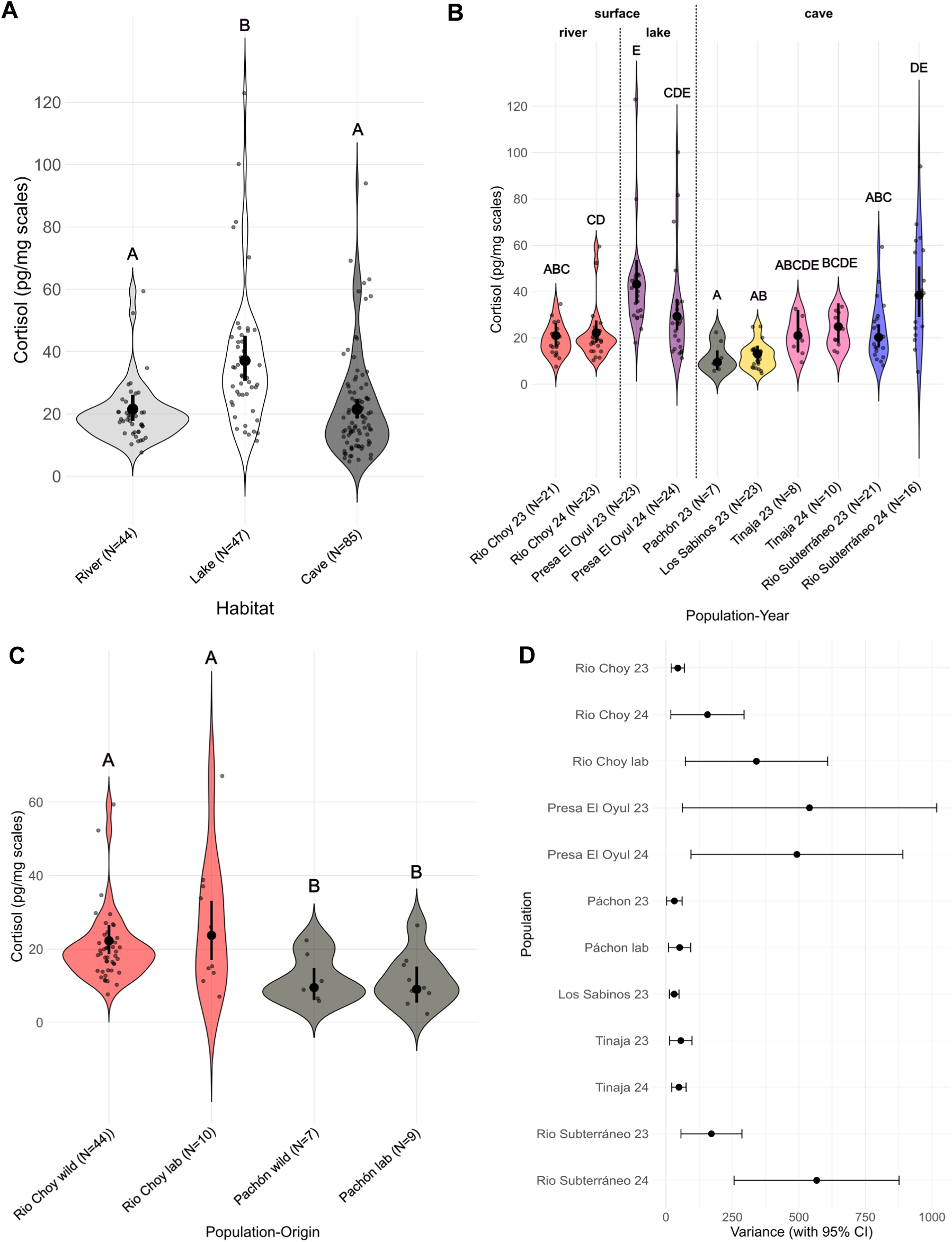
Scale cortisol levels across different habitats and wild and laboratory populations of *A. mexicanus*. (A) Scale cortisol levels in pg/mg scales of wild river (Rio Choy) and lake (Presa El Oyul) surface fish *vs.* cavefish (Pachón, Los Sabinos, Tinaja, Rio Subterráneo) with respective sample size (N) in brackets. The small points and violins show the distribution of the raw data. Large points with error bars show the back-transformed (from log to normal) estimated marginal means (EMMs) with 95% confidence intervals (CIs) under model 1 with scale cortisol as response variable, habitat (lake, river and cave) as predictor, and body condition (K-index) and sex as fixed effects. Significant differences (adjusted p-value <0.05) are indicated by the compact letter design with two habitats being significantly different when no letter overlaps. (B) Scale cortisol levels in pg/mg scales of wild surface and cave populations with respective sample size (N) in brackets. The small points and violins show the distribution of the raw data. Large points with error bars show the back-transformed EMMs with 95% CIs under model 2 with scale cortisol as response variable, population × year as predictor, and body condition (K-index) and sex as fixed effects. Significant differences (adjusted p-value <0.05) are indicated by the compact letter design with two populations × years being significantly different when no letter overlaps. (C) Scale cortisol levels in pg/mg scales of wild *vs.* laboratory Rio Choy surface and Pachón cavefish with respective sample size (N) in brackets. The small points and violins show the distribution of the raw data. Large points with error bars show the back-transformed EMMs with 95% CIs under model 3 with scale cortisol as response variable, population × origin (laboratory vs. wild) as predictor, and body condition (K-index) and sex as fixed effects. Significant differences (adjusted p-value <0.05) are indicated by the compact letter design with two populations × years being significantly different when no letter overlaps. (D) Individual variation in scale cortisol levels displayed by 95% confidence intervals (error bars) with means (points) based on bootstrap resampling (1000 iterations) per population and year/origin (laboratory *vs.* wild).

Comparing populations and years with model 2 revealed that Pachón cavefish in 2023 had the lowest scale cortisol levels (Fig. 2B, see Supplemental Tab. 5 for EMMs), followed by Los Sabinos cavefish. Tinaja cavefish exhibited intermediate cortisol levels in both 2023 and 2024 (Fig. 2B). Interestingly, Rio Choy surface fish showed similar cortisol levels in both wild and laboratory conditions (Fig. 2B). We observed the highest cortisol levels in Presa El Oyul surface fish (Fig. 2B). Rio Subterráneo cavefish exhibited intermediate cortisol levels in 2023, but levels increased in 2024 (Fig. 2B).

### Pairwise comparisons between populations

To assess statistical differences between populations and across years within populations, we conducted pairwise comparisons based on EMMs (see Supplemental Tab. 6 for pairwise comparisons). Pachón and Los Sabinos cavefish exhibited significantly lower scale cortisol levels compared to Rio Choy surface fish in 2024, whereas no significant difference was observed in 2023. However, Pachón cavefish still showed a trend toward lower cortisol levels (Fig. 2B). We observed the largest differences in cortisol levels between Pachón and Los Sabinos cavefish and Presa El Oyul surface fish in both years (Fig. 2B). Comparisons between Tinaja cavefish and both Rio Choy and Presa El Oyul surface fish did not yield any significant differences (Fig.2B). However, Tinaja and Rio Subterráneo cavefish in 2024 exhibited higher cortisol levels than Pachón cavefish (Fig. 2B). We found a notable within-population difference in Rio Subterráneo cavefish, where individuals caught in 2024 had significantly higher cortisol levels than those caught in 2023 (Fig. 2B). Finally, Presa El Oyul surface fish caught in 2023 had significantly higher cortisol levels than Rio Choy surface fish in both years whereas no such difference was observed for Presa El Oyul surface fish in 2024 (Fig. 2B).

### Comparison between laboratory and wild surface and cavefish

In model 3, we tested for differences between wild and laboratory Rio Choy surface and Pachón cavefish with body condition (Fulton’s K-index) and sex as fixed effects. Pairwise comparisons based on EMMs (see Supplemental Tab. 7, 8) revealed that there was neither a significant within ecotype difference in scale cortisol levels between wild and laboratory Rio choy surface fish, nor between wild and laboratory Pachón cavefish (Fig. 2C). However, among ecotypes, Rio Choy surface- and Pachón cavefish differed significantly both between wild and laboratory populations (Fig. 2C).

### Covariate effects on scale cortisol levels

Overall, body condition (Fulton’s k-index) was a significant predictor of cortisol levels in models 1 and 2 and still showed a positive trend in model 3 (Supplemental Tab. 9-11, Supplemental Fig. 4-6), indicating that individuals with better body condition exhibited higher cortisol levels. In contrast, sex did not significantly influence cortisol levels (Supplemental Tab. 9-11, Supplemental Fig. 7-9). In model 2, year had no overall influence on cortisol levels (Supplemental Tab. 10) as well as origin in model 3 (Supplemental Tab. 11).

### Individual variation in scale cortisol levels

To examine individual variation in scale cortisol levels across wild and laboratory populations, we performed bootstrap resampling with 1000 iterations and reported 95% confidence intervals (see Supplemental Tab. 12) for each population × year and population × origin pair. We found that Los Sabinos cavefish in 2023 exhibited the lowest degree of individual variation, followed by wild Rio Choy surface fish in 2023, Tinaja cavefish in 2024, wild Pachón cavefish in 2023, Tinaja cavefish in 2023, and laboratory Pachón cavefish (Fig. 2D). We observed more individual variation in Rio Subterráneo cavefish in 2023, wild Rio Choy surface fish in 2024, laboratory Rio Choy surface fish, and Presa El Oyul surface fish in 2024 (Fig. 2D). Presa El Oyul surface fish in 2023 and Rio Subterráneo cavefish in 2024 (Fig. 2D) exhibited the highest degree of individual variation in scale cortisol levels.

We observed significant differences in individual variation (non-overlap in CIs) between laboratory Rio Choy surface fish and wild Pachón, Los Sabinos, and Tinaja cavefish in 2024 (Fig. 2D). Additionally, Rio Subterráneo cavefish in 2023 and Presa El Oyul surface fish in 2023 showed significantly higher individual variation compared to Los Sabinos cavefish (Fig. 2D). In 2024, both Presa El Oyul surface fish and Rio Subterráneo cavefish exhibited consistently higher individual variation compared to all other cave populations and wild Rio Choy surface fish in 2023 (Fig. 2D).

## Discussion

### Lake habitat effects on scale cortisol levels

Our study provides evidence that long-term experienced stress in *A. mexicanus* varies with habitat type, but not generally between cave and surface ecotypes. Lower scale cortisol levels in cavefish and river surface fish compared to lake surface fish suggest that adaptations to specific habitat characteristics explain variation in scale cortisol levels. Our abiotic measurements indicate that the lake (Presa El Oyul) habitat differed from river surface (Rio Choy) and cave habitats, particularly in temperature, pH, conductivity and iron content. These parameters are known to influence cortisol levels and stress in fish (Choi and An, 2008; Cockrem et al., 2019; Ghazal et al., 2025; Hanke et al., 2019; Lappivaara et al., 1999; Lappivaara and Marttinen, 2005; Mota et al., 2017; O’Toole et al., 2023; Rohner et al., 2013; Samaras et al., 2022; Wagner et al., 1997). However, although differences in these parameters between surface habitats were greater in 2024, we observed no significant difference in scale cortisol levels, suggesting that other environmental factors likely influenced the differences in scale cortisol levels in 2023. Since our abiotic measurements only comprised nine different parameters that were measured once per sampling (and year), they do not represent a dynamic but a snapshot of conditions on the respective day. Hence, more detailed habitat characterizations and long-term tracking of abiotic parameters are needed to better understand the effect of abiotic dynamics on scale cortisol levels in *A. mexicanus*.

Another important aspect to better understand habitat level differences in scale cortisol levels is the time populations could adapt to different habitat types. While river surface and cave populations naturally colonized and adapted to their respective habitat for at least 30,000 to 191,000 years (Fumey et al., 2018; Garduño-Sánchez et al., 2023; Herman et al., 2018; Moran et al., 2023), fish were introduced into the lake Presa El Oyul due to its artificial origin no longer than approximately 45 years ago (“Portal Gob MX,”n.d.). Stressors present in *A. mexicanus* such as predation risk (Chin et al., 2018; Mitchell et al., 1977; Yoshizawa and Library, 2015), aggression (Elipot et al., 2013; Rodriguez-Morales et al., 2022) and parasites (Peuß et al., 2020; Santacruz et al., 2023, 2020) might be different in the natural river and cave vs. artificial lake habitat, which could potentially explain the observed differences in scale cortisol levels between lake and river surface fish and cavefish (Allan et al., 2020; Chin et al., 2018; O’Dwyer et al., 2020; Padmanaban et al., 2025; De Marco et al., 2013; Winberg and Lepage, 1998; Ziv et al., 2012).

### Population differences in scale cortisol levels

Even though we could not show an overall difference in scale cortisol levels between cave and surface fish, we found differences among and within specific cave and surface populations. In 2024, Rio Subterráneo cavefish showed higher scale cortisol levels than Pachón, Los Sabinos and their own population in 2023. Similarly, Tinaja cavefish in 2024 exhibited higher scale cortisol levels than Pachón cavefish. Although our single time-point abiotic measurements showed little variation among these habitats, previous studies have documented seasonal hydrological differences. For instance, unlike Los Sabinos, Tinaja, and Rio Subterráneo, the Pachón cave is not affected by seasonal epigean water inflow due to its elevated entrance (210 m above sea level) (Mitchell et al., 1977). Long-term monitoring confirms that Pachón cave exhibits more stable physico-chemical water parameters, whereas Los Sabinos and Rio Subterráneo experience seasonal fluctuations in temperature, pH, and conductivity (Legendre et al., 2026; Tabin et al., 2018). Thus, combining long-term abiotic data with subsequent scale cortisol sampling could reveal a link between environmental variability and long-term experienced stress.

Differences in the degree of isolation from surface water bodies may also explain within-ecotype and population variation, particularly in Rio Subterráneo. Periodic floodings in this cave increase the influence of seasonal variation, which was shown to impact scale cortisol levels in other fish (Lebigre et al., 2022). In addition, such floodings also introduce surface fish which hybridize with cavefish (Garduño-Sánchez et al., 2023; Moran et al., 2023). Therefore, yearly differences in scale cortisol levels in this population could also result from different sub-samples drawn from a genetically heterogenous cave population (Bradic et al., 2012).

### Genetic determination of scale cortisol levels

We could show that wild Rio Choy surface fish had higher scale cortisol levels than Pachón cavefish. There is direct evidence that Pachón cavefish exhibit altered cortisol regulation compared to Rio Choy surface fish: they show reduced plasma cortisol when exposed to confined space (Gallo and Jeffery, 2012) and lower whole-body cortisol under constant darkness (Bilandžija et al., 2020). Since we did not find significant differences in scale cortisol levels between wild and laboratory-reared Pachón cavefish and Rio Choy surface fish, this suggests that genetic adaptations, rather than plastic responses to short-term environmental conditions, primarily determine long-term stress levels. Furthermore, as the abiotic conditions were different between wild and laboratory populations, this indicates that other genetic adaptations than potential adaptations to the measured parameters might be at play. While cavefish have evolved adaptations that enable them to persist in resource-limited and hypoxic environments over extended periods (Aspiras et al., 2015; Boggs et al., 2022; Olsen et al., 2023; Riddle et al., 2018; Rohner et al., 2013; Shimobayashi et al., 2023; van der Weele and Jeffery, 2022; Xiong et al., 2022, 2018), cortisol responses to food restriction or hypoxia are inconsistent across fish species, being sometimes elevated, reduced or unaffected (Arslan et al., 2016; Caruso et al., 2010; Poursaeid and Falahatkar, 2022; Samaras et al. 2023; Sumpter et al., 1991; White and Fletcher, 1986; Léger et al., 2021; O’Connor et al., 2011). In case of hypoxia, we also observed similarly low scale cortisol levels in Pachón and Los Sabinos cavefish despite strikingly different dissolved oxygen concentrations. Together, this suggests that lower scale cortisol levels in some cave populations might rather be associated with basal metabolic and osmoregulatory processes (Laiz-Carrión et al., 2003; Lawrence et al., 2019) upon food restriction. Therefore, previously discussed stressors as well as basal cortisol regulation in cavefish could be systematically tested in laboratory settings to assess their causal roles in cortisol accumulation in scales. Additionally, genetic mapping (Jeffery, 2020) of scale cortisol levels, through crossbreeding and quantitative trait loci analysis, could help identify underlying genetic loci involved in cortisol regulation. Including more cave populations such as Tinaja and Rio Subterráneo could clarify whether cortisol regulation and potential stress adaptations are convergent or population specific. Furthermore, the functional consequences of reduced cortisol in Pachón cavefish remain complex. While lower cortisol may promote energy allocation to somatic growth (Mommsen et al., 1999), support year-round reproduction (Schreck, 2010; Espinasa et al., 2023; Xia et al., 2025), shift immune investment (Peuß et al., 2020) and contribute to longevity (Cobham et al., 2024; Medley et al., 2022) due to reduced immunosuppression and oxidative stress (Sapolsky et al., 2000), glucocorticoid interactions with life-history traits are multifaceted (Crespi et al., 2013). Sustained cortisol reduction may thus provide important advantages in some cave habitats, but further research is required to clarify these links.

### Individual variation in scale cortisol levels

We observed population-level differences in individual variability of scale cortisol levels. Low individual variation in Los Sabinos, Pachón, and Tinaja cavefish suggests that these cave habitats are characterised by low fluctuations in stressors, leading individuals to occupy similar niches within the given habitat. Adaptations to strong and consistent selective pressures (Poulson, 2001; Venarsky et al., 2014), such as limited food availability and stable, yet sometimes extreme abiotic conditions (Legendre et al., 2025; Poulson and White, 1969; Tabin et al., 2018), may contribute to reduced individual variation in stress responses and long-term stress levels. This reduced variability could, in turn, limit the ability to conform to niche changes through glucocorticoid mediated stress responses. Presa El Oyul Lake potentially exhibits greater environmental heterogeneity and dynamic stressors, including fluctuating abiotic parameters, and potentially, predation, and aggressive interactions among conspecifics, which may explain the higher individual variation in scale cortisol levels. In Rio Subterráneo, periodic flooding from nearby surface streams, along with the introduction and hybridization of surface individuals (Garduño-Sánchez et al., 2023; Moran et al., 2023) together with invasive parasites (Santacruz et al., 2023), could have contributed to greater individual variation in scale cortisol levels. This variation may stem from gradients of cave-adapted traits across the sampled individuals leading to differences in stress responses.

In general, there might also be differences in the inter-individual cortisol responsiveness among populations. A recent study found that individuals responding to stress with either high or low cortisol levels can also be identified based on their scale cortisol levels (Samaras et al., 2021) which could explain the observed differences in individual variation.

### Scale cortisol as a measure for long-term stress

Surprisingly, we found a significant positive relationship between body condition (K-index) and scale cortisol levels, contrasting with previous studies in other fish species, where higher scale cortisol levels were either associated with reduced body condition (Brosset et al., 2023) or showed no correlation (Carbajal et al., 2019). One possible explanation is that cavefish exhibit increased fat accumulation (Xiong et al., 2022, 2018), which elevates Fulton’s K-index through increased body mass. In humans, obesity and metabolic syndrome are linked to increased cortisol levels and hyperactivity of the hypothalamic-pituitary-adrenal axis (Anagnostis et al., 2009). A similar mechanism may influence cortisol regulation in *A. mexicanus*, warranting further research into interactions between cortisol, fat storage, and body condition in this species.

Even though scales have been shown to be capable of releasing cortisol in cell culture medium upon stimulation (Samaras and Pavlidis, 2022), it is unlikely that potential short-term released cortisol due to capture and handling of fish accumulated in scale tissue in such a short amount of time, as it usually takes longer than 72h after an acute stressor to elevate scale cortisol levels (Laberge et al., 2019). Therefore, our scale cortisol levels likely reflect long-term cortisol accumulation. Nonetheless, future studies should aim to integrate precise age estimates, such as counting annual growth increments on scales or otoliths (Simon et al., 2017), into scale cortisol analyses of wild populations and directly test whether cortisol accumulates in scales over time in *A. mexicanus* to support its suitability as a measure for long-term-experienced stress.

While we also cannot rule out a possible bias introduced by differential catching methods (dip-netting vs. angling), the use of both methods was necessary to ensure adequate sampling across habitats. Both angling and dip-netting can produce non-random samples of fish populations such as selecting for bolder or larger/smaller individuals (Ghazal et al., 2025; Thorburn, 1992) with potential consequences for scale cortisol levels. Therefore, both methods should be further tested with respect to scale cortisol sampling in *A. mexicanus*.

Furthermore, the actual responsiveness to cortisol needs to be investigated in this system since there could be individual or population differences due to varying glucocorticoid receptor expression, as shown for example in arctic charr and trout (Johansen et al., 2011; Vindas et al., 2017), which can be organ and time-specific (Teles et al., 2013). Teleost fish also have different glucocorticoid receptor variants with different sensitivities and metabolic functions (Stolte et al., 2006). It is also important to consider that an individual’s stress phenotype cannot be determined solely by cortisol levels, as both high and low levels may be either pathological or adaptive (Romero and Beattie, 2022). Therefore, it is necessary to include other measures of stress such as catecholamines (Reid et al., 1998) and stress-induced behaviours (Balasch and Tort, 2019) into an integrated view on the stress response in *A. mexicanus*.

## Conclusion

Our findings suggest that long-term stress, as indicated by scale cortisol levels, is higher in lake surface fish compared to river surface and cavefish, potentially due to specific habitat characteristics and adaptations. Furthermore, we could show that there are differences in scale cortisol levels among cave populations and between river surface fish and the most isolated cave population. Genetic adaptations to habitat specific stressors appear to shape long-term stress responses across populations, emphasizing the role of stress physiology in colonizing diverse and extreme habitat types. Future research should experimentally test the effects of specific stressors, such as hypoxia, food restriction, abiotic parameters, predation, parasites and aggression on cortisol regulation and explore the genetic basis of stress-related differences between ecotypes. Our study highlights the importance of scale cortisol as a biomarker for long-term stress and provides insights into how habitat and population specific adaptations shape long-term stress.

## Competing interests

The authors declare no competing interests.

### Acknowledgements

We thank Michelle Borgers, Ulla Pebesma, Annika Potthoff, David Perez Guerra, Riley Kellermeyer for their help with fieldwork and fish dissections. We also acknowledge people from Ejido Los Sabinos and from the Comisión Nacional de Áreas Naturales Protegidas (CONANP) for their guidance in Los Sabinos and Tinaja caves. Furthermore, we thank Edith Ossendorf and Sabine Kruse for support on establishing and conducting the ELISA assay. Finally, we thank Jaime Anaya-Rojas for advice with statistical analysis and Nicolas Rohner for critical comments on a previous version of this manuscript. This work was funded by the German Research Foundation (DFG) as part of the CRC TRR 212 (NC3, 316099922), projects B06 (granted to R.P., Project number 471673683) and project S, Endocrinology platform (granted to S.K., Project number 396782989).

## Author contributions

**Marc Bauhus:** investigation (lead); formal analysis (lead); project administration (lead); writing – original draft (lead); visualization (lead); conceptualization (support). **Luise Wander**: investigation (support); formal analysis (support); writing – review & editing (equal). **David Boy:** investigation (support); formal analysis (support); writing – review & editing (equal). **Ana Santacruz:** investigation (support); writing – review & editing (equal). **Ernesto Maldonado:** investigation (support); writing – review & editing (equal). **Joachim Kurtz:** resources (support); supervision (support); writing – original draft (support); writing – review & editing (equal). **Sylvia Kaiser:** methodology (lead); resources (support); funding acquisition (support); writing – review & editing (equal). **Robert Peuß:** conceptualization (lead); funding acquisition (lead); project administration (support); supervision (lead); resources (lead); visualization (support); writing – original draft (support); writing – review & editing (equal).

## Statement on inclusion

Our study brings together authors from a number of different countries, including scientists based in the country where the study was carried out. All authors were engaged early on with the research and study design to ensure that the diverse sets of perspectives they represent was considered from the onset. Whenever possible, our research was discussed with the local community to inform on our science and the consequences that our field work potentially has on the local ecosystem.

## Ethics statement

All wild animals were collected under permit no. SPARN/DGVS/03371/23 granted by the Secretaría de Medio Ambiente y Recursos Naturales (SEMARNAT) to Ernesto Maldonado. All animals were treated in accordance with local veterinary and animal welfare authorities and their respective guidelines. Scales from laboratory fish were used under the permit T24.016UMS granted by the University of Münster, Germany in accordance with local veterinary and animal welfare authorities.

## Data availability statement

Data and code can be found under https://github.com/Marc9704/Amex_scale_cortisolG

## References

Aerts J, Metz JR, Ampe B, Decostere A, Flik G, De Saeger S. 2015. Scales Tell a Story on the Stress History of Fish. PLoS One 10:e0123411. doi:10.1371/journal.pone.0123411

Allan BJM, Illing BR, Fakan EP, Narvaez P, Sgrutter A, Sikkel PC, McClure EC, Rummer JL, McCormick MI. 2020. Parasite infection directly impacts escape response and stress levels in fish. Journal of Experimental Biology 223. doi:10.1242/jeb.230904

Anagnostis P, Athyros VG, Tziomalos K, Karagiannis A, Mikhailidis DP. 2009. The Pathogenetic Role of Cortisol in the Metabolic Syndrome: A Hypothesis. J Clin Endocrinol Metab 94:2692–2701. doi:10.1210/JC.2009-0370

Arslan G, Sahin T, Hisar O, Hisar SA. 2016. Effects of low temperature and starvation on plasma cortisol, triiodothyronine, thyroxine, thyroid-stimulating hormone and prolactin levels of juvenile common carp (*Cyprinus carpio*). Mar Sci Technol Bull 4:5–9.

Aspiras AC, Rohner N, Martineau B, Borowsky RL, Tabin CJ. 2015. Melanocortin 4 receptor mutations contribute to the adaptation of cavefish to nutrient-poor conditions. Proc Natl Acad Sci U S A 112:9668–9673. doi:10.1073/pnas.1510802112

Balasch JC, Tort L. 2019. Netting the stress responses in fish. Front Endocrinol (Lausanne) 10:435714. doi:10.3389/fendo.2019.00062

Barton B, Morgan J, Vijayan M. 2002. Physiological and condition-related indicators of environmental stress in fish.

Baumann DP, Ingalls A. 2022. Mexican tetra (*Astyanax mexicanus*): biology, husbandry, and experimental protocols. Laboratory Fish in Biomedical Research: Biology, Husbandry and Research Applications for Zebrafish, Medaka, Killifish, Cavefish, Stickleback, Goldfish and Danionella Translucida 311–347. doi:10.1016/B978-0-12-821099-4.00003-1

Bilandžija H, Hollifield B, Steck M, Meng G, Ng M, Koch AD, Gračan R, Ćetković H, Porter ML, Renner KJ, Jeffery W. 2020. Phenotypic plasticity as a mechanism of cave colonization and adaptation. Elife 9. doi:10.7554/elife.51830

Boggs TE, Friedman JS, Gross JB. 2022. Alterations to cavefish red blood cells provide evidence of adaptation to reduced subterranean oxygen. Scientific Reports 2022 12:1 12:1–10. doi:10.1038/s41598-022-07619-0

Bradic M, Beerli P, García-De Leán FJ, Esquivel-Bobadilla S, Borowsky RL. 2012. Gene flow and population structure in the Mexican blind cavefish complex (*Astyanax mexicanus*). BMC Evol Biol 12:1–17. doi:10.1186/1471-2148-12-9

Brillon DJ, Zheng B, Campbell RG, Matthews DE. 1995. Effect of cortisol on energy expenditure and amino acid metabolism in humans. https://doi.org/101152/ajpendo19952683E501 268. doi:10.1152/ajpendo.1995.268.3.E501

Britton JR, Andreou D, Lopez-Bejar M, Carbajal A. 2023. Relationships of scale cortisol content suggest stress resilience in freshwater fish vulnerable to catch-and-release angling in recreational fisheries. Fish Res 266:106776. doi:10.1016/j.fishres.2023.106776

Brosset P, Averty A, Mathieu-Resuge M, Schull Q, Soudant P, Lebigre C. 2023. Fish morphometric body condition indices reflect energy reserves but other physiological processes matter. Ecol Indic 154:110860. doi:10.1016/j.ecolind.2023.110860

Carbajal A, Monclús L, Tallo-Parra O, Sabes-Alsina M, Vinyoles D, Lopez-Bejar M. 2018. Cortisol detection in fish scales by enzyme immunoassay: Biochemical and methodological validation. Journal of Applied Ichthyology 34:967–970. doi:10.1111/jai.13674

Carbajal A, Tallo-Parra O, Monclús L, Vinyoles D, Solé M, Lacorte S, Lopez-Bejar M. 2019. Variation in scale cortisol concentrations of a wild freshwater fish: Habitat quality or seasonal influences? Gen Comp Endocrinol 275:44–50. doi:10.1016/j.ygcen.2019.01.015

Caruso G, Maricchiolo G, Micale V, Genovese L, Caruso R, Denaro MG. 2010. Physiological responses to starvation in the European eel (Anguilla anguilla): Effects on haematological, biochemical, non-specific immune parameters and skin structures. Fish Physiol Biochem 36:71–83. doi:10.1007/S10695-008-9290-6

Chin JSR, Gassant CE, Amaral PM, Lloyd E, Stahl BA, Jaggard JB, Keene AC, Duboue ER. 2018. Convergence on reduced stress behavior in the Mexican blind cavefish. Dev Biol 441:319–327. doi:10.1016/j.ydbio.2018.05.009

Choi CY, An KW. 2008. Cloning and expression of Na+/K+-ATPase and osmotic stress transcription factor 1 mRNA in black porgy, *Acanthopagrus schlegeli* during osmotic stress. Comp Biochem Physiol B Biochem Mol Biol 149:91–100. doi:10.1016/j.cbpb.2007.08.009

Christiansen JJ, Djurhuus CB, Gravholt CH, Iversen P, Christiansen JS, Schmitz O, Weeke J, Jørgensen JOL, Møller N. 2007. Effects of Cortisol on Carbohydrate, Lipid, and Protein Metabolism: Studies of Acute Cortisol Withdrawal in Adrenocortical Failure. J Clin Endocrinol Metab 92:3553–3559. doi:10.1210/jc.2007-0445

Cobham AE, Kenzior A, Morales-Sosa P, Javier JE, Swanson S, Wood C, Rohner N. 2024. Cave Adaptation Favors Aging Resilience in the Mexican Tetra. bioRxiv 2024.09.26.615235. doi:10.1101/2024.09.26.615235

Cockrem JF, Bahry MA, Chowdhury VS. 2019. Cortisol responses of goldfish (Carassius auratus) to air exposure, chasing, and increased water temperature. Gen Comp Endocrinol 270:18–25. doi:10.1016/j.ygcen.2018.09.017

Crespi EJ, Williams TD, Jessop TS, Delehanty B. 2013. Life history and the ecology of stress: How do glucocorticoid hormones influence life-history variation in animals? Funct Ecol 27:93–106. doi:10.1111/1365-2435.12009

De Marco RJ, Groneberg AH, Yeh CM, Castillo Ramírez LA, Ryu S. 2013. Optogenetic elevation of endogenous glucocorticoid level in larval zebrafish. Front Neural Circuits 7:46978. doi:10.3389/fncir.2013.00082

DeKoning ABL, Picard DJ, Bond SR, Schulte PM. 2004. Stress and interpopulation variation in glycolytic enzyme activity and expression in a teleost fish *Fundulus heteroclitus*. Physiological and Biochemical Zoology 77:18–26. doi:10.1086/378914

Ding J, Fanjara E, Stene A, Gansel LC, Finstad B, Cao Y. 2024. Comparison and Validation of Commercial Elisa Kits for Quantification of Fecal Cortisol Metabolites in Atlantic Salmon (Salmon Salar). doi:10.2139/ssrn.5012212

Elipot Y, Hinaux H, Callebert J, Rétaux S. 2013. Evolutionary shift from fighting to foraging in blind cavefish through changes in the serotonin network. Current Biology 23:1–10. doi:10.1016/j.cub.2012.10.044

Espinasa L, Rohner N, Rétaux S. 2023. Reproductive seasonality of *Astyanax mexicanus* cavefish. Zool Res 44:698. doi:10.24272/j.issn.2095-8137.2022.164

Fumey J, Hinaux H, Noirot C, Thermes C, Rétaux S, Casane D. 2018. Evidence for late Pleistocene origin of Astyanax mexicanus cavefish. BMC Evolutionary Biology 18:1–19. DOI: 10.1186/S12862-018-1156-7

Gallo ND, Jeffery WR. 2012. Evolution of Space Dependent Growth in the Teleost *Astyanax mexicanus*. PLoS One 7:e41443. doi:10.1371/journal.pone.0041443

Garduño-Sánchez M, Hernández-Lozano J, Moran RL, Miranda-Gamboa R, Gross JB, Rohner N, Elliott WR, Miller J, Lozano-Vilano L, McGaugh SE, Ornelas-García CP. 2023. Phylogeographic relationships and morphological evolution between cave and surface *Astyanax mexicanus* populations (De Filippi 1853) (Actinopterygii, Characidae). Mol Ecol 32:5626–5644. doi:10.1111/MEC.17128

Gates RD, Baghdasarian G, Muscatine L. 1992. Temperature Stress Causes Host Cell Detachment in Symbiotic Cnidarians: Implications for Coral Bleaching. 182:324–332. doi:10.2307/1542252

Ghazal A, Paul R, Tarkan AS, Britton JR. 2025. Influence of season, capture method, sample age and extraction protocols on the scale cortisol concentrations of three species of freshwater fish. General and Comparative Endocrinology 362:114671. DOI: 10.1016/j.ygcen.2025.114671

Hanke I, Ampe B, Kunzmann A, Gärdes A, Aerts J. 2019. Thermal stress response of juvenile milkfish (Chanos chanos) quantified by ontogenetic and regenerated scale cortisol. Aquaculture 500:24–30. DOI: 10.1016/j.aquaculture.2018.09.016

Herman A, Brandvain Y, Weagley J, Jeffery WR, Keene AC, Kono TJY, Bilandžija H, Borowsky R, Espinasa L, O’Quin K, Ornelas-García CP, Yoshizawa M, Carlson B, Maldonado E, Gross JB, Cartwright RA, Rohner N, Warren WC, McGaugh SE. 2018. The role of gene flow in rapid and repeated evolution of cave-related traits in Mexican tetra, Astyanax mexicanus. Molecular Ecology 27:4397–4416. DOI: 10.1111/mec.14877

Huffman DL, Abrami L, Sasik R, Corbeil J, Van Der Goot FG, Aroian R V. 2004. Mitogen-activated protein kinase pathways defend against bacterial pore-forming toxins. Proc Natl Acad Sci U S A 101:10995–11000. doi:10.1073/pnas.0404073101

Jeffery WR. 2020. *Astyanax* surface and cave fish morphs. Evodevo 11:1–10. doi:10.1186/S13227-020-00159-6

Johansen IB, Sandvik GK, Nilsson GE, Bakken M, Øverli Ø. 2011. Cortisol receptor expression differs in the brains of rainbow trout selected for divergent cortisol responses. Comp Biochem Physiol Part D Genomics Proteomics 6:126–132. doi:10.1016/j.cbd.2010.11.002

Krishnan J, Persons JL, Peuß R, Hassan H, Kenzior A, Xiong S, Olsen L, Maldonado E, Kowalko JE, Rohner N. 2020. Comparative transcriptome analysis of wild and lab populations of *Astyanax mexicanus* uncovers differential effects of environment and morphotype on gene expression. J Exp Zool B Mol Dev Evol. doi:10.1002/jez.b.22933

Laberge F, Yin-Liao I, Bernier NJ. 2019. Temporal profiles of cortisol accumulation and clearance support scale cortisol content as an indicator of chronic stress in fish. Conserv Physiol 7:621–626. doi:10.1093/conphys/coz052

Laiz-Carrión R, Martín Del Río MP, Miguez JM, Mangera JM, Soengas JL. 2003. Influence of cortisol on osmoregulation and energy metabolism in gilthead seabream Sparus aurata. Journal of Experimental Zoology Part A: Comparative Experimental Biology 298A:105–118. DOI: 10.1002/jez.a.10256

Lappivaara J, Kiviniemi A, Oikari A. 1999. Bioaccumulation and subchronic physiological effects of waterborne iron overload on whitefish exposed in humic and nonhumic water. Arch Environ Contam Toxicol 37:196–204. doi:10.1007/S002449900506

Lappivaara J, Marttinen S. 2005. Effects of waterborne iron overload and simulated winter conditions on acute physiological stress response of whitefish, *Coregonus lavaretus*. Ecotoxicol Environ Saf 60:157–168. doi:10.1016/j.ecoenv.2004.01.003

Lawrence MJ, Eliason EJ, Zolderdo AJ, Lapointe D, Best C, Gilmour KM, Cooke SJ. 2019. Cortisol modulates metabolism and energy mobilization in wild-caught pumpkinseed (Lepomis gibbosus). Fish Physiology and Biochemistry 45:1813–1828. DOI: 10.1007/S10695-019-00680-Z

Lebigre C, Woillez M, Barone H, Mourot J, Drogou M, Le Goff R, Servili A, Hennebert J, Vanhomwegen M, Aerts J. 2022. Temporal variations in scale cortisol indicate consistent local-and broad-scale constraints in a wild marine teleost fish. Mar Environ Res 182:105783. doi:10.1016/j.marenvres.2022.105783

Legendre L, Père S, Rebaudo F, Espinasa L, Attia J, Rétaux S. 2026. Water Parameters and Hydrodynamics in Rivers and Caves Hosting Astyanax mexicanus Populations Reveal Macro-, Meso- and Microhabitat Characteristics. Ecology and Evolution 16:e72970. DOI: 10.1002/ECE3.72970

Léger JAD, Athanasio CG, Zhera A, Chauhan MF, Simmons DBD. 2021. Hypoxic responses in Oncorhynchus mykiss involve angiogenesis, lipid, and lactate metabolism, which may be triggered by the cortisol stress response and epigenetic methylation. Comp Biochem Physiol Part D Genomics Proteomics 39:100860. doi:10.1016/j.cbd.2021.100860

Medley JK, Persons J, Biswas T, Olsen L, Peuß R, Krishnan J, Xiong S, Rohner N. 2022. The metabolome of Mexican cavefish shows a convergent signature highlighting sugar, antioxidant, and Ageing-Related metabolites. Elife 11. doi:10.7554/elife.74539

Mitchell R, Russell W, Elliott W. 1977. Mexican eyeless Characin fishes, genus *Astyanax*: environment, distribution, and evolution. KIP Monographs.

Mommsen TP, Vijayan MM, Moon TW. 1999. Cortisol in teleosts: Dynamics, mechanisms of action, and metabolic regulation. Rev Fish Biol Fish 9:211–268. doi:10.1023/A:1008924418720

Moran RL, Richards EJ, Ornelas-García CP, Gross JB, Donny A, Wiese J, Keene AC, Kowalko JE, Rohner N, McGaugh SE. 2023. Selection-driven trait loss in independently evolved cavefish populations. Nature Communications 2023 14:1 14:1–19. doi:10.1038/s41467-023-37909-8

Mota VC, Martins CIM, Eding EH, Canário AVM, Verreth JAJ. 2017. Cortisol and testosterone accumulation in a low pH recirculating aquaculture system for rainbow trout (*Oncorhynchus mykiss*). Aquac Res 48:3579–3588. doi:10.1111/are.13184

Näslund J, Rosengren M, Del Villar D, Gansel L, Norrgård JR, Persson L, Winkowski JJ, Kvingedal E. 2013. Hatchery tank enrichment affects cortisol levels and shelter-seeking in Atlantic salmon (Salmo salar). Canadian Journal of Fisheries and Aquatic Sciences 70:585–590. DOI: 10.1139/cjfas-2012-0302

O’Connor EA, Pottinger TG, Sneddon LU. 2011. The effects of acute and chronic hypoxia on cortisol, glucose and lactate concentrations in different populations of three-spined stickleback. Fish Physiol Biochem 37:461–469. doi:10.1007/S10695-010-9447-Y

O’Dwyer K, Dargent F, Forbes MR, Koprivnikar J. 2020. Parasite infection leads to widespread glucocorticoid hormone increases in vertebrate hosts: A meta-analysis. Journal of Animal Ecology 89:519–529. doi:10.1111/1365-2656.13123

O’Toole C, White P, Thomas K, O’Maoiléidigh N, Fjelldal PG, Hansen TJ, Graham CT, Brophy D. 2023. Effects of temperature and feeding regime on cortisol concentrations in scales of Atlantic salmon post-smolts. Journal of Experimental Marine Biology and Ecology 569:151955. DOI: 10.1016/j.jembe.2023.151955

Olsen L, Levy M, Medley JK, Hassan H, Miller B, Alexander R, Wilcock E, Yi K, Florens L, Weaver K, McKinney SA, Peuß R, Persons J, Kenzior A, Maldonado E, Delventhal K, Gluesenkamp A, Mager E, Coughlin D, Rohner N. 2023. Metabolic reprogramming underlies cavefish muscular endurance despite loss of muscle mass and contractility. Proc Natl Acad Sci U S A 120:e2204427120. doi:10.1073/pnas.2204427120

Ornelas-García P, Pajares S, Sosa-Jiménez VM, Rétaux S, Miranda-Gamboa RA. 2018. Microbiome differences between river-dwelling and cave-adapted populations of the fish *Astyanax mexicanus* (De Filippi, 1853). PeerJ 2018:e5906. doi:10.7717/peerj.5906

Padmanaban N, Ambosie R, Choy S, Marcus S, Nilsson SRO, Keene AC, Kowalko JE, Duboué ER. 2025. Automated Behavioral Profiling Using Neural Networks Reveals Differences in Stress-Like Behavior Between Cave and Surface-Dwelling *Astyanax mexicanus*. J Exp Zool B Mol Dev Evol 344:352–362. doi:10.1002/jez.b.23311

Peuß R, Box AC, Chen S, Wang Y, Tsuchiya D, Persons JL, Kenzior A, Maldonado E, Krishnan J, Scharsack JP, Slaughter BD, Rohner N. 2020. Adaptation to low parasite abundance affects immune investment and immunopathological responses of cavefish. Nat Ecol Evol. doi:10.1038/s41559-020-1234-2

Portal Gob MX. n.d. https://www.gob.mx/

Poulson TL. 2001. Adaptations of cave fishes with some comparisons to deep-sea fishes. Environ Biol Fishes 62:345–364. doi:10.1023/A:1011893916855

Poulson TL, White WB. 1969. The cave environment. Science (1979) 165:971–981. doi:10.1126/science.165.3897.971

Poursaeid S, Falahatkar B. 2022. Starvation alters growth, stress metabolites and physiological responses in juvenile great sturgeon (*Huso huso*). Anim Feed Sci Technol 294:115429. doi:10.1016/j.anifeedsci.2022.115429

Pravosudov V V., Kitaysky AS, Wingfield JC, Clayton NS. 2001. Long-Term Unpredictable Foraging Conditions and Physiological Stress Response in Mountain Chickadees (*Poecile gambeli*). Gen Comp Endocrinol 123:324–331. doi:10.1006/gcen.2001.7684

Reid SG, Bernier NJ, Perry SF. 1998. The adrenergic stress response in fish: control of catecholamine storage and release. Comp Biochem Physiol C Pharmacol Toxicol Endocrinol 120:1–27. doi:10.1016/S0742-8413(98)00037-1

Riddle MR, Aspiras AC, Gaudenz K, Peuß R, Sung JY, Martineau B, Peavey M, Box AC, Tabin JA, McGaugh S, Borowsky R, Tabin CJ, Rohner N. 2018. Insulin resistance in cavefish as an adaptation to a nutrient-limited environment. Nature 2018 555:7698 555:647–651. doi:10.1038/nature26136

Rodriguez-Morales R, Gonzalez-Lerma P, Yuiska A, Han JH, Guerra Y, Crisostomo L, Keene AC, Duboue ER, Kowalko JE. 2022. Convergence on reduced aggression through shared behavioral traits in multiple populations of *Astyanax mexicanus*. BMC Ecol Evol 22:1–15. doi:10.1186/S12862-022-02069-8

Rohner N, Jarosz DF, Kowalko JE, Yoshizawa M, Jeffery WR, Borowsky RL, Lindquist S, Tabin CJ. 2013. Cryptic variation in morphological evolution: HSP90 as a capacitor for loss of eyes in cavefish. Science (1979) 342:1372–1375. doi:10.1126/science.1240276

Romero LM, Beattie UK. 2022. Common myths of glucocorticoid function in ecology and conservation. J Exp Zool A Ecol Integr Physiol 337:7–14. doi:10.1002/jez.2459

Roque d’orbcastel E, Bettarel Y, Dellinger M, Sadoul B, Bouvier T, Amandé JM, Dagorn L, Geffroy B. 2021. Measuring cortisol in fish scales to study stress in wild tropical tuna. Environ Biol Fishes 104:725–732. doi:10.1007/S10641-021-01107-6

Sachser N, Lick C. 1991. Social experience, behavior, and stress in guinea pigs. Physiol Behav 50:83–90. doi:10.1016/0031-9384(91)90502-F

Samaras A, Dimitroglou A, Kollias S, Skouradakis G, Papadakis IE, Pavlidis M. 2021. Cortisol concentration in scales is a valid indicator for the assessment of chronic stress in European sea bass, Dicentrarchus labrax L. Aquaculture 545:737257. DOI: 10.1016/j.aquaculture.2021.737257

Samaras A, Dimitroglou A, Gleni KE, Pavlidis M. 2022. Physiological responses of red seabream (Pagrus major) to stress and rearing temperature. Aquaculture Research 53:2518–2528. DOI: 10.1111/are.15771

Samaras A, Pavlidis M. 2022. Fish Scales Produce Cortisol upon Stimulation with ACTH. Animals 2022, Vol 12, Page 3510 12:3510. doi:10.3390/ani12243510

Samaras A, Tsoukali P, Katsika L, Pavlidis M, Papadakis IE. 2023. Chronic impact of exposure to low dissolved oxygen on the physiology of Dicentrarchus labrax and Sparus aurata and its effects on the acute stress response. Aquaculture 562:738830. DOI: 10.1016/j.aquaculture.2022.738830

Santacruz A, Claudia |, Ornelas-García P, Pérez-Ponce De León G, Mérida U, Mérida M. 2020. Incipient genetic divergence or cryptic speciation? *Procamallanus* (Nematoda) in freshwater fishes (*Astyanax*). Zool Scr 49:768–778. doi:10.1111/zsc.12443

Santacruz A, Hernández-Mena D, Miranda-Gamboa R, De León GPP, Ornelas-García CP. 2023. Host-parasite interactions in perpetual darkness: Macroparasite diversity in the cavefish *Astyanax mexicanus*. Zool Res 44:782. doi:10.24272/j.issn.2095-8137.2022.376

Sapolsky RM, Romero LM, Munck AU. 2000. How Do Glucocorticoids Influence Stress Responses? Integrating Permissive, Suppressive, Stimulatory, and Preparative Actions. Endocr Rev 21:55–89. doi:10.1210/edrv.21.1.0389

Schreck CB. 2010. Stress and fish reproduction: The roles of allostasis and hormesis. Gen Comp Endocrinol 165:549–556. doi:10.1016/j.ygcen.2009.07.004

Schulte PM, Davies SA, Dow JAT, Lukowiak K. 2014. What is environmental stress? Insights from fish living in a variable environment. Journal of Experimental Biology 217:23–34. doi:10.1242/jeb.089722

Shimobayashi M, Thomas A, Shetty S, Frei IC, Wölnerhanssen BK, Weissenberger D, Vandekeere A, Planque M, Dietz N, Ritz D, Meyer-Gerspach AC, Maier T, Hay N, Peterli R, Fendt SM, Rohner N, Hall MN. 2023. Diet-induced loss of adipose hexokinase 2 correlates with hyperglycemia. Elife 12. doi:10.7554/elife.85103

Simon V, Elleboode R, Mahé K, Legendre L, Ornelas-Garcia P, Espinasa L, Rétaux S. 2017. Comparing growth in surface and cave morphs of the species Astyanax mexicanus: insights from scales. SpringerV Simon, R Elleboode, K Mahé, L Legendre, P Ornelas-Garcia, L Espinasa, S RétauxEvodevo, 2017•Springer 8. doi:10.1186/s13227-017-0086-6

Stolte EH, Verburg van Kemenade BML, Savelkoul HFJ, Flik G. 2006. Evolution of glucocorticoid receptors with different glucocorticoid sensitivity. Journal of Endocrinology 190:17–28. doi:10.1677/joe.1.06703

Sumpter JP, Le Bail PY, Pickering AD, Pottinger TG, Carragher JF. 1991. The effect of starvation on growth and plasma growth hormone concentrations of rainbow trout, *Oncorhynchus mykiss*. Gen Comp Endocrinol 83:94–102. doi:10.1016/0016-6480(91)90109-J

Tabin JA, Aspiras A, Martineau B, Riddle M, Kowalko J, Borowsky R, Rohner N, Tabin CJ. 2018. Temperature preference of cave and surface populations of *Astyanax mexicanus*. Dev Biol 441:338–344. doi:10.1016/j.ydbio.2018.04.017

Teles M, Tridico R, Callol A, Fierro-Castro C, Tort L. 2013. Differential expression of the corticosteroid receptors GR1, GR2 and MR in rainbow trout organs with slow release cortisol implants. Comp Biochem Physiol A Mol Integr Physiol 164:506–511. doi:10.1016/j.cbpa.2012.12.018

Thorburn MA. 1992. The randomness of samples collected by dip-net methods from rainbow trout in tanks. Aquaculture 101:385–390. DOI: 10.1016/0044-8486(92)90041-I

van der Weele CM, Jeffery WR. 2022. Cavefish cope with environmental hypoxia by developing more erythrocytes and overexpression of hypoxia-inducible genes. Elife 11. doi:10.7554/elife.69109

Van Dievel M, Janssens L, Stoks R. 2016. Short- and long-term behavioural, physiological and stoichiometric responses to predation risk indicate chronic stress and compensatory mechanisms. Oecologia 181:347–357. doi:10.1007/S00442-015-3440-1

Van Raaij MTM, Pit DSS, Balm PHM, Steffens AB, Van Den Thillart GEEJM. 1996. Behavioral Strategy and the Physiological Stress Response in Rainbow Trout Exposed to Severe Hypoxia. Horm Behav 30:85–92. doi:10.1006/hbeh.1996.0012

Venarsky MP, Huntsman BM, Huryn AD, Benstead JP, Kuhajda BR. 2014. Quantitative food web analysis supports the energy-limitation hypothesis in cave stream ecosystems. Oecologia 176:859–869. doi:10.1007/S00442-014-3042-3

Vercauteren M, Ampe B, Devriese L, Moons CPH, Decostere A, Aerts J, Chiers K. 2022. Explorative study on scale cortisol accumulation in wild caught common dab (*Limanda limanda*). BMC Vet Res 18:1–12. doi:10.1186/S12917-022-03385-3

Vijayan MM, Pereira C, Grau EG, Iwama GK. 1997. Metabolic Responses Associated with Confinement Stress in Tilapia: The Role of Cortisol. Comp Biochem Physiol C Pharmacol Toxicol Endocrinol 116:89–95. doi:10.1016/S0742-8413(96)00124-7

Vindas MA, Magnhagen C, Brännäs E, Øverli Ø, Winberg S, Nilsson J, Backström T. 2017. Brain cortisol receptor expression differs in Arctic charr displaying opposite coping styles. Physiol Behav 177:161–168. doi:10.1016/j.physbeh.2017.04.024

Wagner EJ, Bosakowski,’ And T, Intelmann S. 1997. Combined Effects of Temperature and High pH on Mortality and the Stress Response of Rainbow Trout after Stocking; Combined Effects of Temperature and High pH on Mortality and the Stress Response of Rainbow Trout after Stocking. doi:10.1577/1548-8659(1997)126

Wang WN, Zhou J, Wang P, Tian TT, Zheng Y, Liu Y, Mai W jun, Wang AL. 2009. Oxidative stress, DNA damage and antioxidant enzyme gene expression in the Pacific white shrimp, *Litopenaeus vannamei* when exposed to acute pH stress. Comparative Biochemistry and Physiology Part C: Toxicology & Pharmacology 150:428–435. doi:10.1016/j.cbpc.2009.06.010

Webster Marketon JI, Glaser R. 2008. Stress hormones and immune function. Cell Immunol 252:16–26. doi:10.1016/j.cellimm.2007.09.006

White A, Fletcher TC. 1986. Serum cortisol, glucose and lipids in plaice (*Pleuronectes platessa L.*) exposed to starvation and aquarium stress. Comp Biochem Physiol A Physiol 84:649–653. doi:10.1016/0300-9629(86)90380-4

Winberg S, Lepage O. 1998. Elevation of brain 5-HT activity, POMC expression, and plasma cortisol in socially subordinate rainbow trout. Am J Physiol Regul Integr Comp Physiol 274. doi:10.1152/ajpregu.1998.274.3.R645

Xia F, Santacruz A, Wu D, Bertho S, Fritz E, Morales-Sosa P, McKinney S, Nowotarski SH, Rohner N. 2025. Reproductive adaptation of *Astyanax mexicanus* under nutrient limitation. Dev Biol 523:82–98. doi:10.1016/j.ydbio.2025.04.006

Xiong S, Krishnan J, Peuß R, Rohner N. 2018. Early adipogenesis contributes to excess fat accumulation in cave populations of *Astyanax mexicanus*. Dev Biol 441:297–304. doi:10.1016/j.ydbio.2018.06.003

Xiong S, Wang W, Kenzior A, Olsen L, Krishnan J, Persons J, Medley K, Peuß R, Wang Y, Chen S, Zhang N, Thomas N, Miles JM, Alvarado AS, Rohner N. 2022. Enhanced lipogenesis through Pparγ helps cavefish adapt to food scarcity. Current Biology 32:2272–2280.e6. doi:10.1016/j.cub.2022.03.038

Yoshizawa M, Library WO. 2015. Behaviors of cavefish offer insight into developmental evolution. Mol Reprod Dev 82:268–280. doi:10.1002/mrd.22471

Yoshizawa M, Robinson BG, Duboué ER, Masek P, Jaggard JB, O’Quin KE, Borowsky RL, Jeffery WR, Keene AC. 2015. Distinct genetic architecture underlies the emergence of sleep loss and prey-seeking behavior in the Mexican cavefish. BMC Biol 13:1–12. doi:10.1186/S12915-015-0119-3

Zhang Z, Xu X, Wang Y, Zhang X. 2020. Effects of environmental enrichment on growth performance, aggressive behavior and stress-induced changes in cortisol release and neurogenesis of black rockfish Sebastes schlegelii. Aquaculture 528:735483. DOI: 10.1016/j.aquaculture.2020.735483

Ziv L, Muto A, Schoonheim PJ, Meijsing SH, Strasser D, Ingraham HA, Schaaf MJM, Yamamoto KR, Baier H. 2012. An affective disorder in zebrafish with mutation of the glucocorticoid receptor. Molecular Psychiatry 2013 18:6 18:681–691. doi:10.1038/mp.2012.64

