## Supplemental Information for "Scale cortisol levels vary with habitat type and population in the Mexican tetra *Astyanax mexicanus*"

#### Methods - Assay quality control

##### Inter- and intra-assay variations and cross-reactions

Three control samples were used to determine the inter-assay variation. They were determined in duplicate on a total of six plates. The inter-assay deviation was 8.79 on average. Intra-assay coefficients of variation (CV) were calculated from all duplicated samples analysed. It was on average 3.96. Finally, all samples were assayed in duplicate and the determination of the sample was repeated if the CV was larger than 10%. The antibody showed the following cross-reactivities: Cortisol 100.0%, Prednisolone 47.4%, Cortisone 15.7%, 11-Deoxycortisol 15.0%, Prednisone 7.83%, Corticosterone 4.81%, 6β-Hydroxycortisol 1.37%, 17-Hydroxyprogesterone 1.36%, Deoxycorticosterone 0.94%, Progesterone 0.06%, Bethamethasone 0.05%, Dehydroepiandrosterone 0.03%, Dexamethasone 0.03%.

**Table 1. Linearity** To assess linearity, we ran a serial dilution of three pooled samples in duplicate.

| | expected (pg/ml) | measured (pg/ml) | linearity (%) |
|---|---|---|---|
| **sample 1** | | | |
| neat | | 2153 | |
| 1:2 | 1077 | 988 | **92** |
| 1:4 | 538 | 525 | **98** |
| 1:8 | 269 | 280 | **104** |
| 1:16 | 134 | 134 | **100** |
| **sample 2** | | | |
| neat | | 1190 | |
| 1:2 | 595 | 600 | **100.8** |
| 1:4 | 298 | 270 | **90.8** |
| 1:8 | 149 | 130 | **87.4** |
| **sample 3** | | | |
| neat | | 3160 | |
| 1:2 | 1580 | 1470 | **93.0** |
| 1:4 | 790 | 650 | **82.3** |
| 1:8 | 395 | 316 | **80.0** |
| 1:16 | 198 | 150 | **75.8** |

**Recovery rate**

Three pooled samples of different concentrations were used to calculate the recovery rate of the assay. We conducted spikes by mixing equal volumes of the respective pooled sample with three different standards, which were also used for the standard curve in the ELISA. All samples were assayed in duplicate. The expected recovery concentrations were based on the known amount of cortisol concentrations in the control samples.

Table 2. Recovery rate: To assess the recovery rate, three pooled samples of different concentrations were spiked with different amounts of corticosterone and measured in duplicate.

| | expected (pg/ml) | measured (pg/ml) | recovery rate (%) |
|---|---|---|---|
| **sample 1** | | | |
| Initial value | | 480 | **100.0** |
| + 400 pg/ml | 680 (480+200) | 650 | **95.6** |
| + 1000 pg/ml | 980 (480+500) | 980 | **93.9** |
| + 2000 pg/ml | 1480 (480+1000) | 1480 | **85.5** |
| **Sample 2** | | | |
| Initial value | | 380 | **100.0** |
| + 400 pg/ml | 580 (380+200) | 510 | **87.9** |
| + 1000 pg/ml | 880 (380+500) | 730 | **83.0** |
| + 2000 pg/ml | 1380 (380+1000) | 1210 | **87.7** |
| **Sample 3** | | | |
| Initial value | | 230 | **100.0** |
| + 400 pg/ml | 430 (230+200) | 400 | **93.0** |
| + 1000 pg/ml | 730 (230+500) | 710 | **97.3** |
| + 2000 pg/ml | 1230 (230+1000) | 1060 | **86.2** |

#### Figures

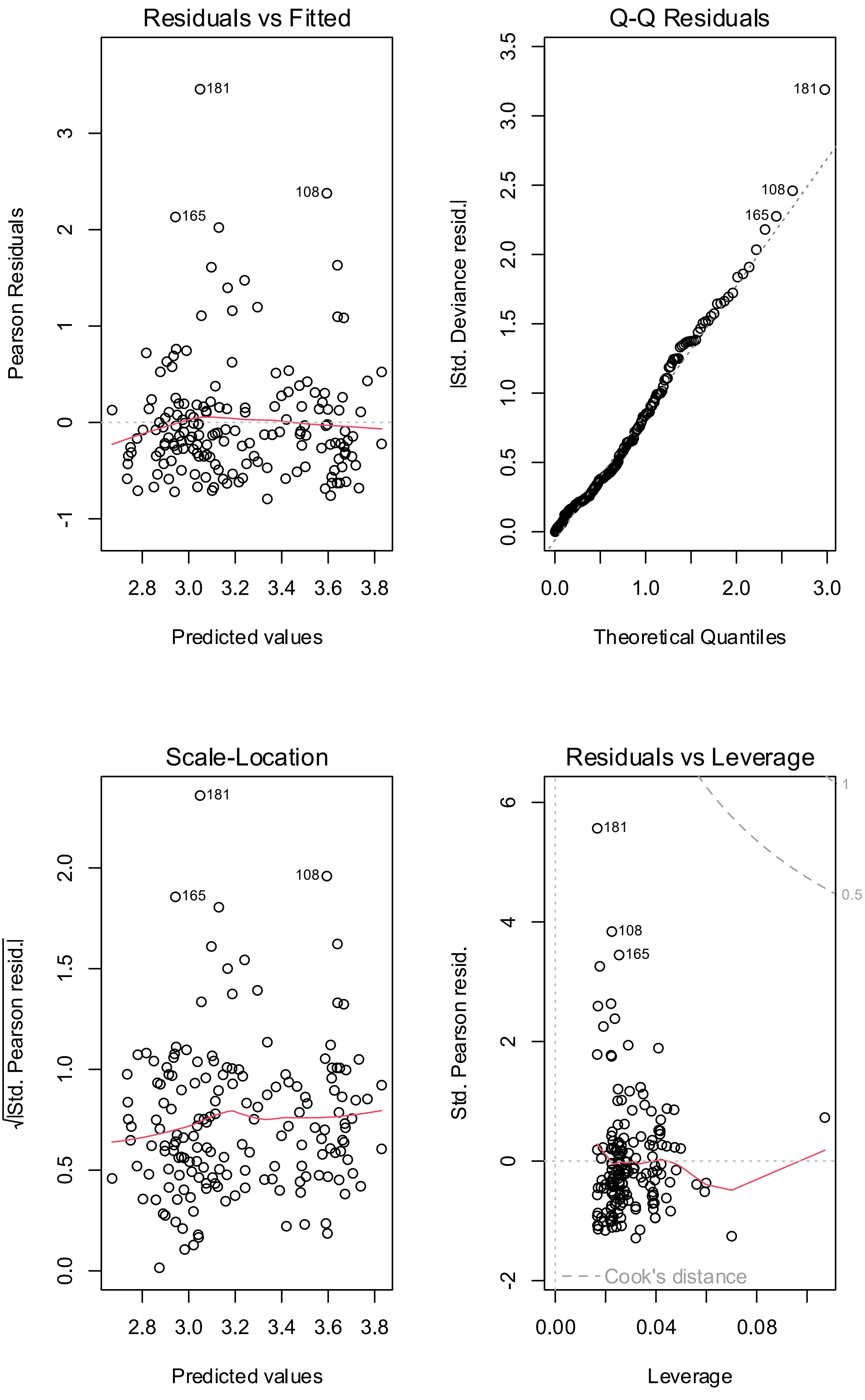

Residuals vs Fitted
Pearson Residuals
Predicted values
181
108
165
Q-Q Residuals
|Std. Deviance resid.|
Theoretical Quantiles
181
108
165
Scale-Location
√|Std. Pearson resid.|
Predicted values
181
108
165
Residuals vs Leverage
Std. Pearson resid.
Leverage
181
108
165
Cook's distance
1
0.5

**Supplemental Figure 1.** Model diagnostic plots for model 1 testing for an association of scale cortisol levels (response variable) and habitat (predictor) with body condition (Fulton's K index) and sex as coviates. (A) Residuals vs. Fitted, assessing non-linearity and variance patterns. The y-axis represents the residuals (e.g., Pearson or deviance residuals), while the x-axis shows the fitted values. The red smoothing line visualizes potential trends deviating from the dotted reference line at zero; (B) Normal Q-Q, checking the normality of deviance residuals. The x-axis represents the expected quantiles from a normal distribution, while the y-axis shows actual quantiles of the residuals. The dotted diagonal reference line represents a normal distribution; (C) Scale-Location, evaluating homoscedasticity. The y-axis represents the square root of the absolute standardized residuals, and the x-axis shows the fitted values. The red smoothing line visualizes systematic trends; and (D) Residuals vs. Leverage, detecting influential observations. The x-axis represents leverage, indicating how much an observation influences model estimates. The y-axis shows standardized residuals, with dashed Cook's distance contours helping identify highly influential points. The red smoothing line provides a visual indication of the overall trend in residuals.

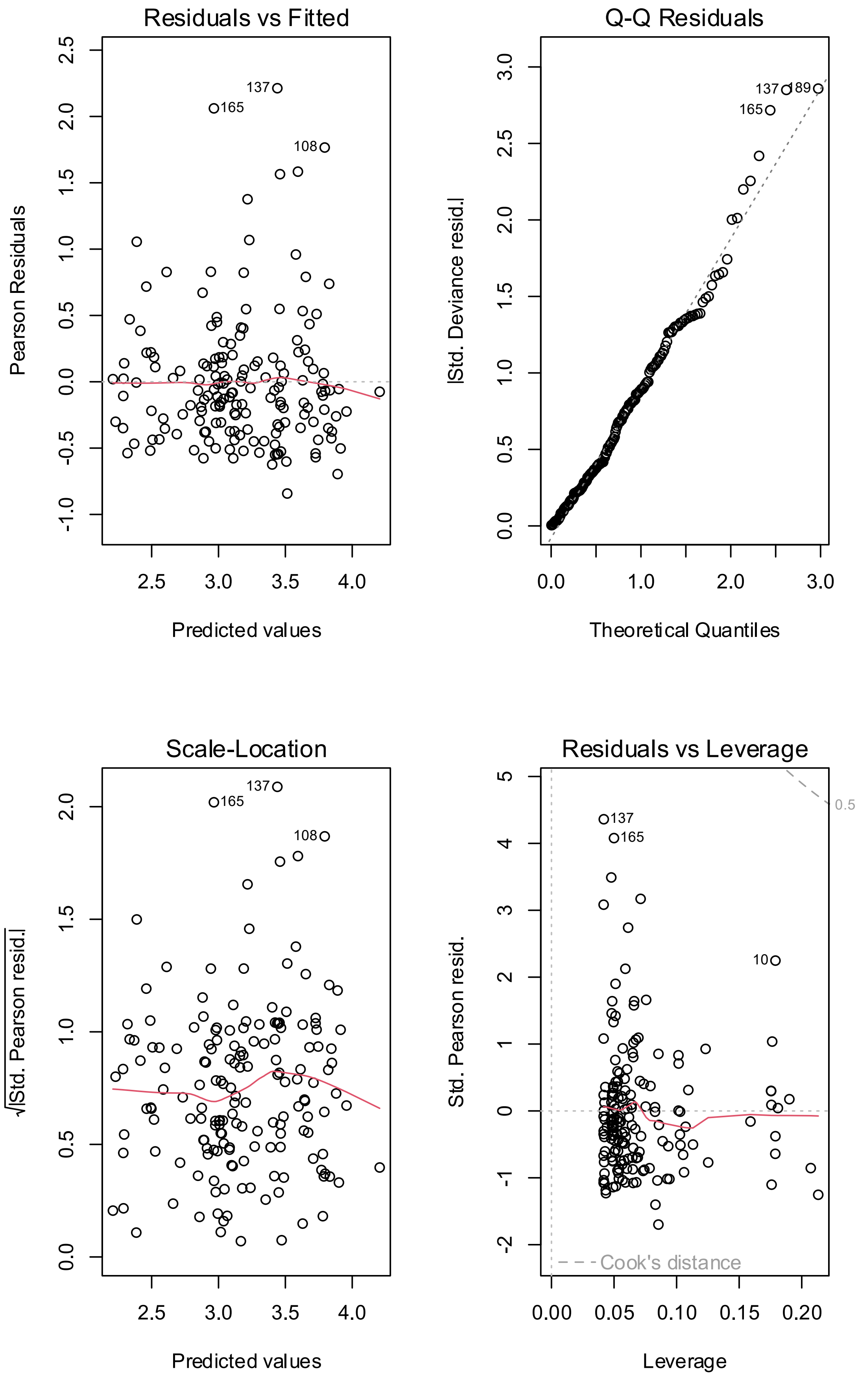

Residuals vs Fitted
Pearson Residuals
Predicted values
137
165
108
Q-Q Residuals
|Std. Deviance resid.|
Theoretical Quantiles
137
189
165
Scale-Location
√|Std. Pearson resid.|
Predicted values
137
165
108
Residuals vs Leverage
Std. Pearson resid.
Leverage
Cook's distance
0.5
137
165
10

**Supplemental Figure 2.** Model diagnostic plots for model 2 testing for an association of scale cortisol levels (response variable) and population x year with body condition (Fulton's K index) and sex as coviates. (A) Residuals vs. Fitted, assessing non-linearity and variance patterns. The y-axis represents the residuals (e.g., Pearson or deviance residuals), while the x-axis shows the fitted values. The red smoothing line visualizes potential trends deviating from the dotted reference line at zero; (B) Normal Q-Q, checking the normality of deviance residuals. The x-axis represents the expected quantiles from a normal distribution, while the y-axis shows actual quantiles of the residuals. The dotted diagonal reference line represents a normal distribution; (C) Scale-Location, evaluating homoscedasticity. The y-axis represents the square root of the absolute standardized residuals, and the x-axis shows the fitted values. The red smoothing line visualizes systematic trends; and (D) Residuals vs. Leverage, detecting influential observations. The x-axis represents leverage, indicating how much an observation influences model estimates. The y-axis shows standardized residuals, with dashed Cook's distance contours helping identify highly influential points. The red smoothing line provides a visual indication of the overall trend in residuals.

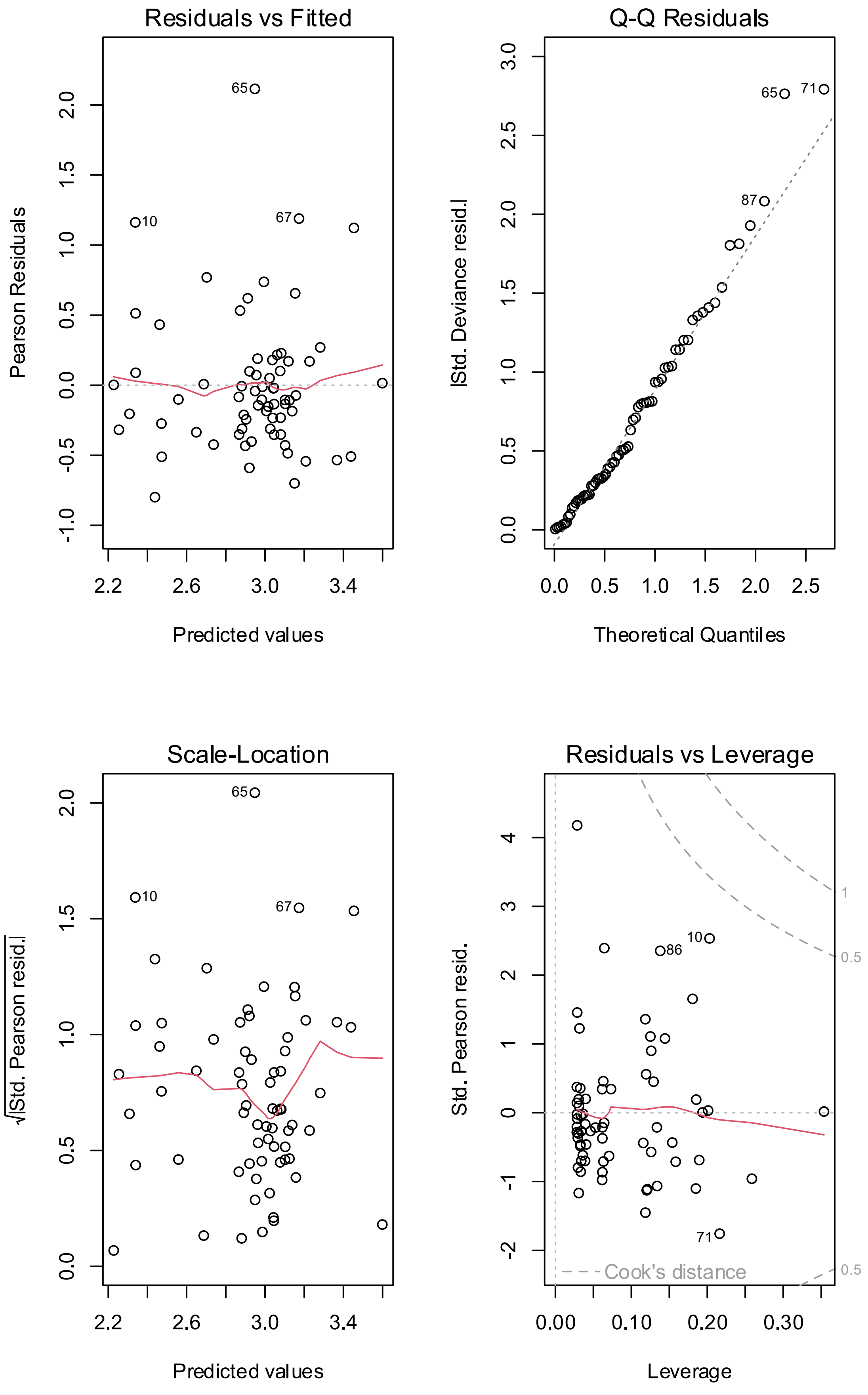

Residuals vs Fitted
Pearson Residuals
Predicted values
Q-Q Residuals
|Std. Deviance resid.|
Theoretical Quantiles
Scale-Location
√|Std. Pearson resid.|
Predicted values
Residuals vs Leverage
Std. Pearson resid.
Leverage
Cook's distance

**Supplemental Figure 3.** Model diagnostic plots for model 3 testing for an association of scale cortisol levels (response variable) and population x origin with body condition (Fulton's K index) and sex as coviates. (A) Residuals vs. Fitted, assessing non-linearity and variance patterns. The y-axis represents the residuals (e.g., Pearson or deviance residuals), while the x-axis shows the fitted values. The red smoothing line visualizes potential trends deviating from the dotted reference line at zero; (B) Normal Q-Q, checking the normality of deviance residuals. The x-axis represents the expected quantiles from a normal distribution, while the y-axis shows actual quantiles of the residuals. The dotted diagonal reference line represents a normal distribution; (C) Scale-Location, evaluating homoscedasticity. The y-axis represents the square root of the absolute standardized residuals, and the x-axis shows the fitted values. The red smoothing line visualizes systematic trends; and (D) Residuals vs. Leverage, detecting influential observations. The x-axis represents leverage, indicating how much an observation influences model estimates. The y-axis shows standardized residuals, with dashed Cook's distance contours helping identify highly influential points. The red smoothing line provides a visual indication of the overall trend in residuals.

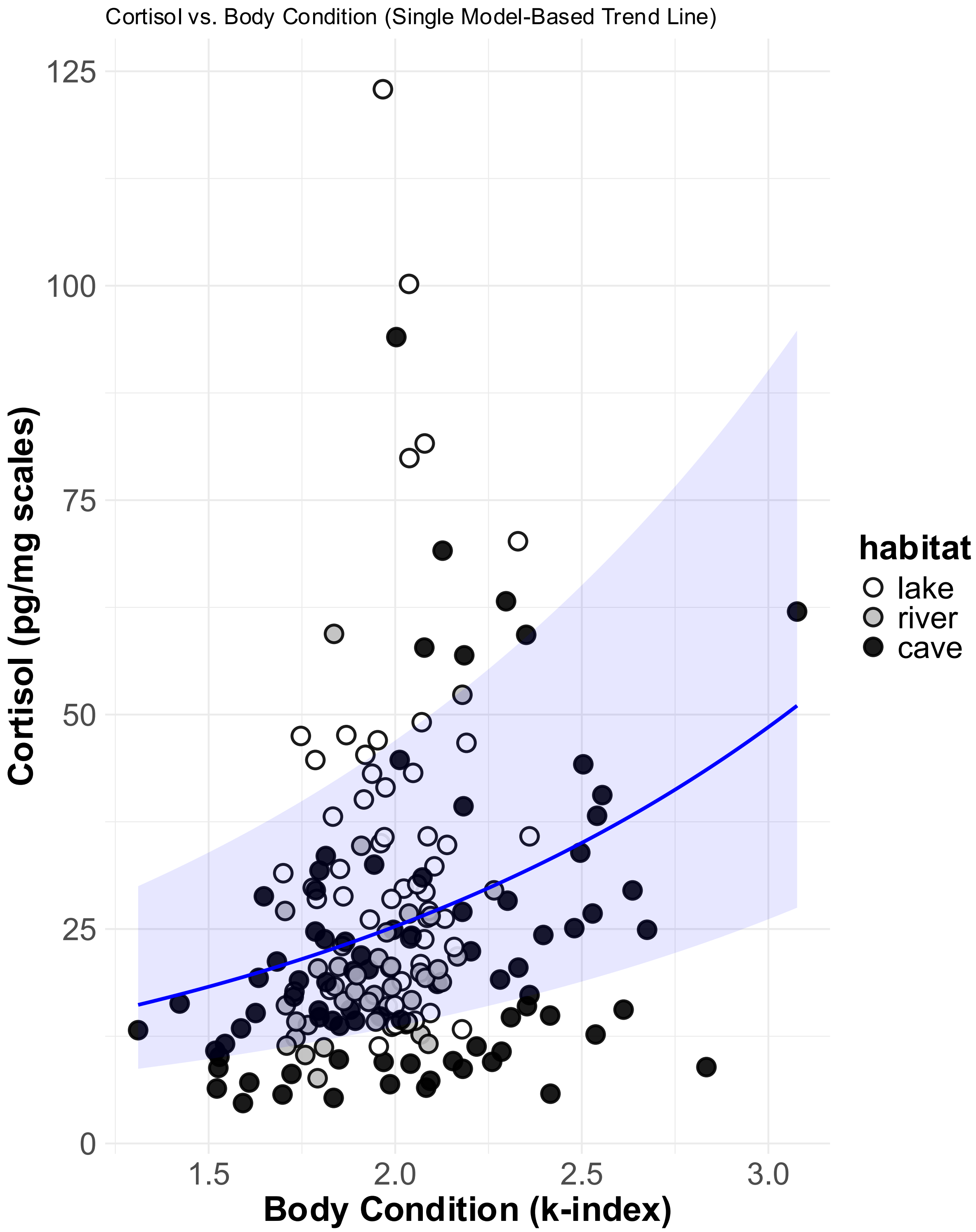

**Supplemental Figure 4.** Correlation between body condition (Fulton's k-index) and scale cortisol levels with raw data points colored by habitat. The blue trend line represents predictions based on model 1 averaged across habitat and the shaded area indicates the 95% confidence intervals around the trend line.

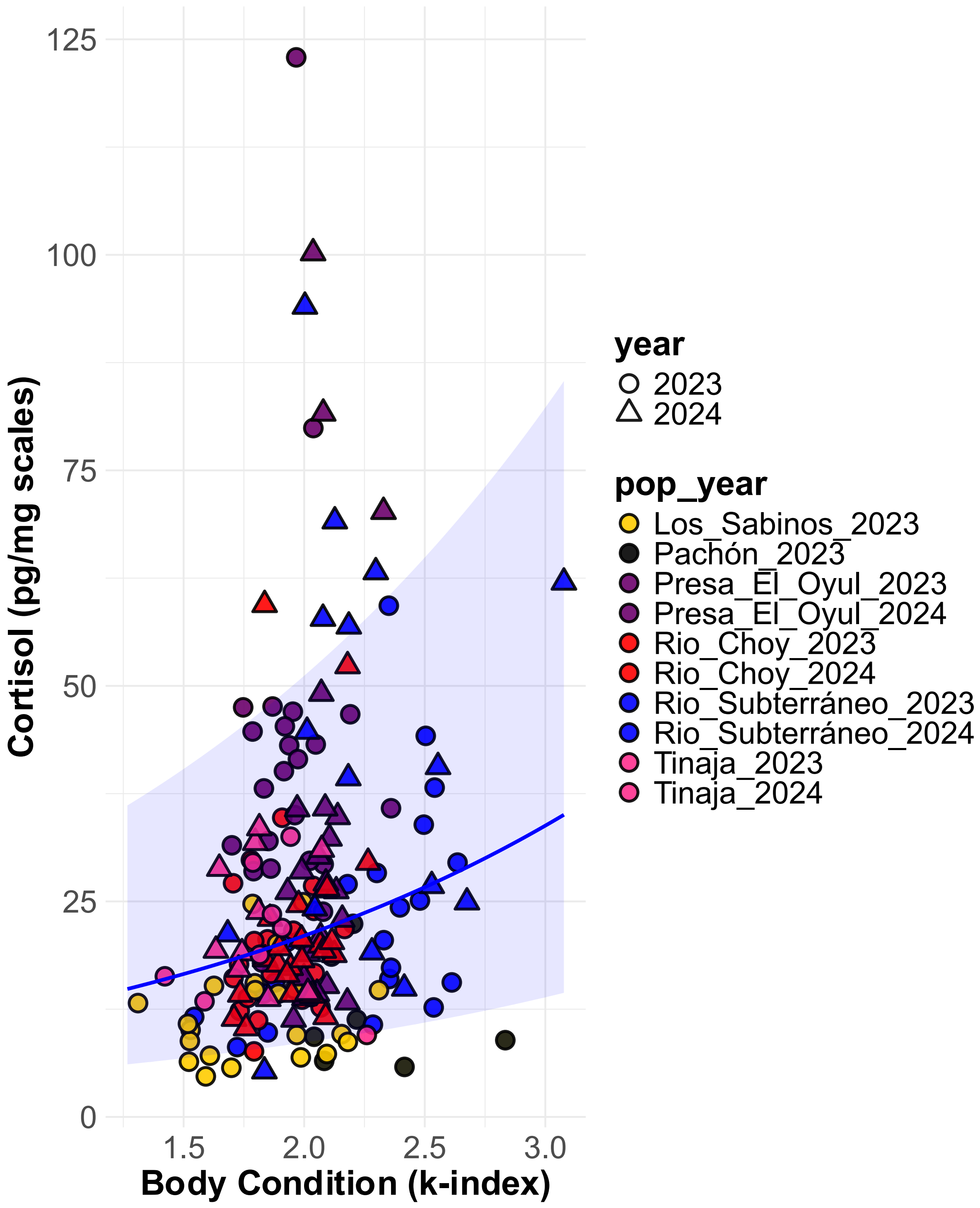

**Supplemental Figure 5.** Correlation between body condition (Fulton's k-index) and scale cortisol levels with raw data points colored by population × year (pop_year). The blue trend line represents predictions based on model 2 averaged across population × year and the shaded area indicates the 95% confidence intervals around the trend line.

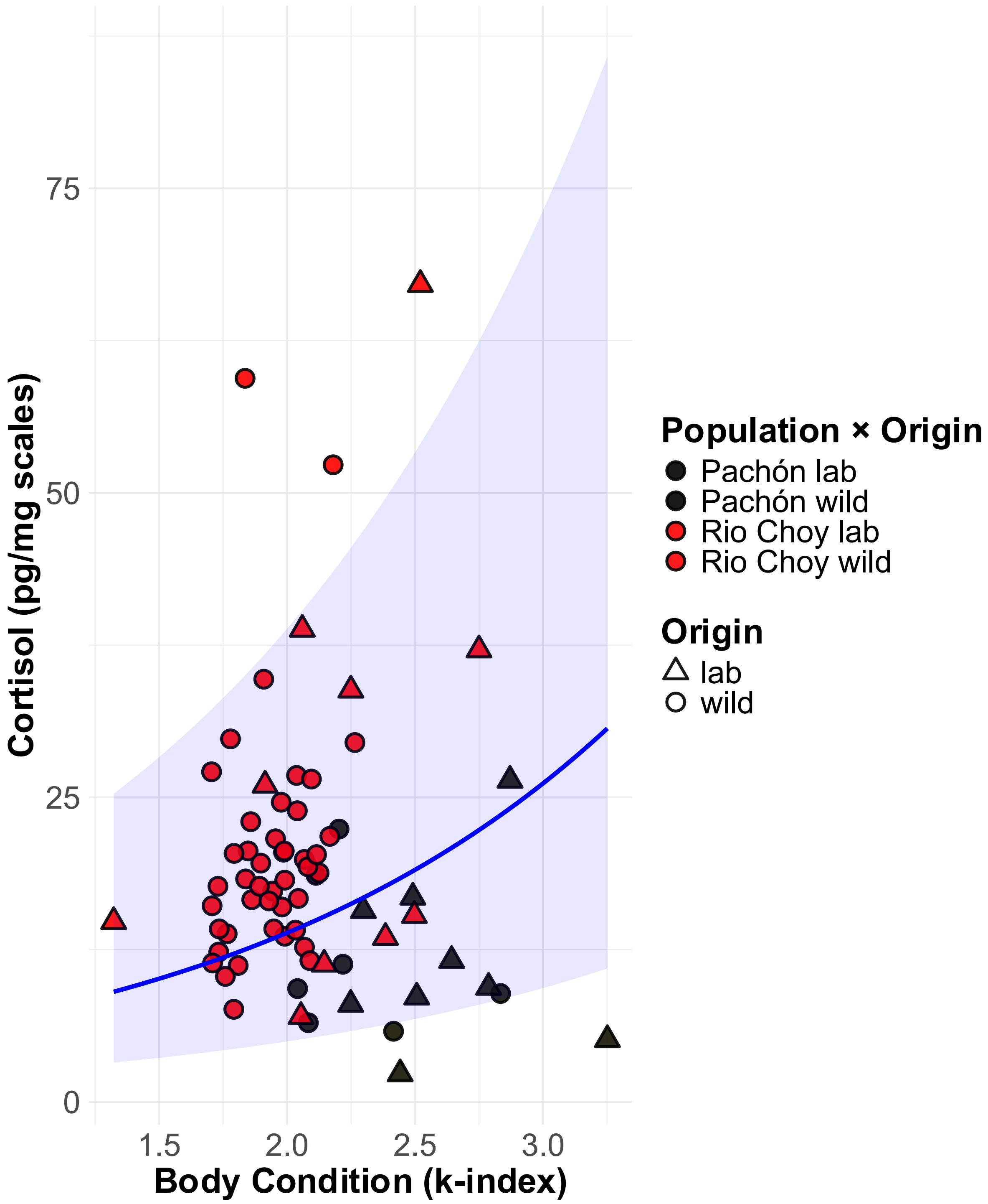

**Supplemental Figure 6.** Correlation between body condition (Fulton's k-index) and scale cortisol levels with raw data points colored by population × origin. The blue trend line represents predictions based on model 3 averaged across population × origin and the shaded area indicates the 95% confidence intervals around the trend line.

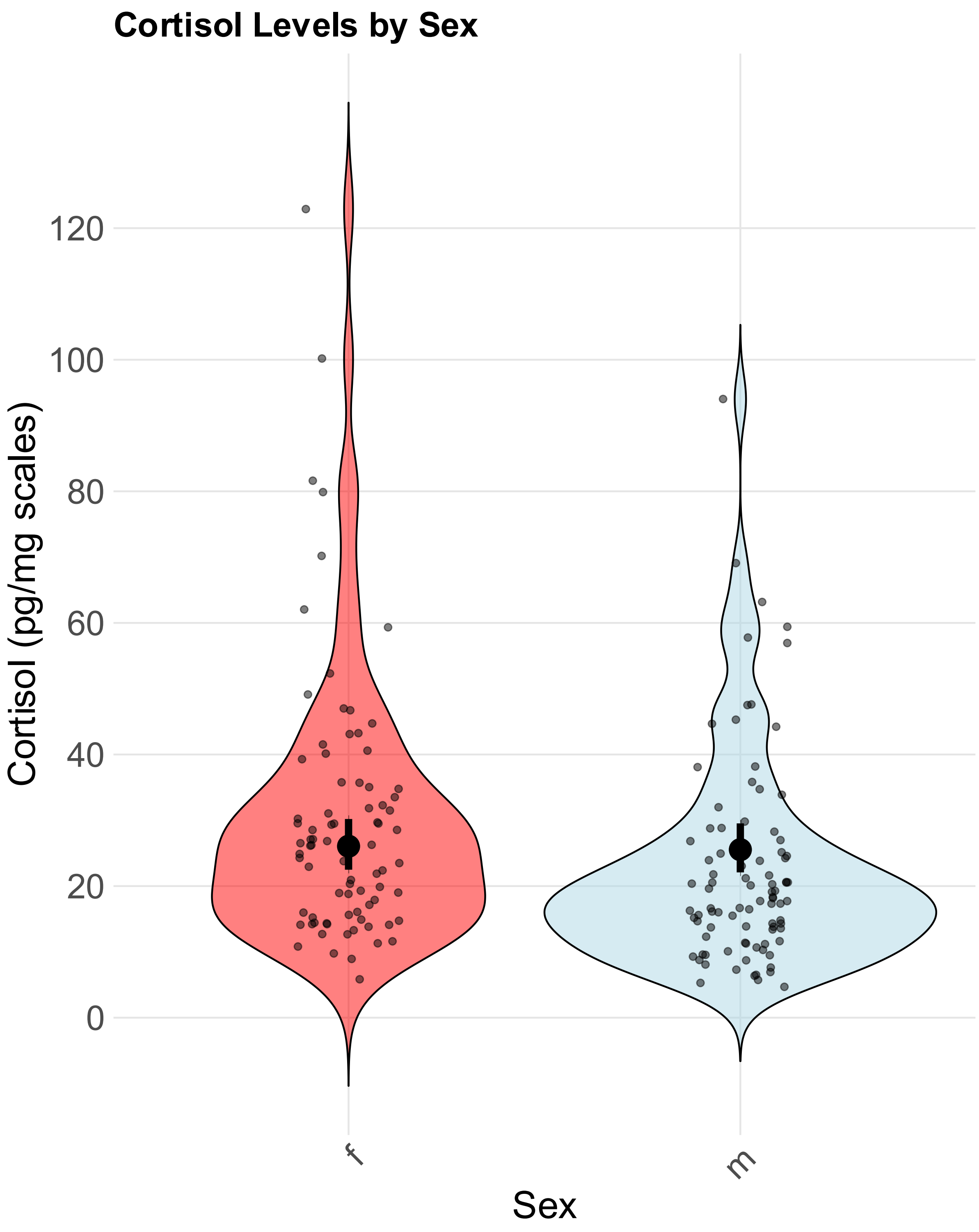

**Supplemental Figure 7.** Cortisol levels by sex based on model 1 testing for habitat differences. Large points with error bars represent the estimated marginal mean from the model for females (f) and males (m). Light points and violins show the distribution of the raw data.

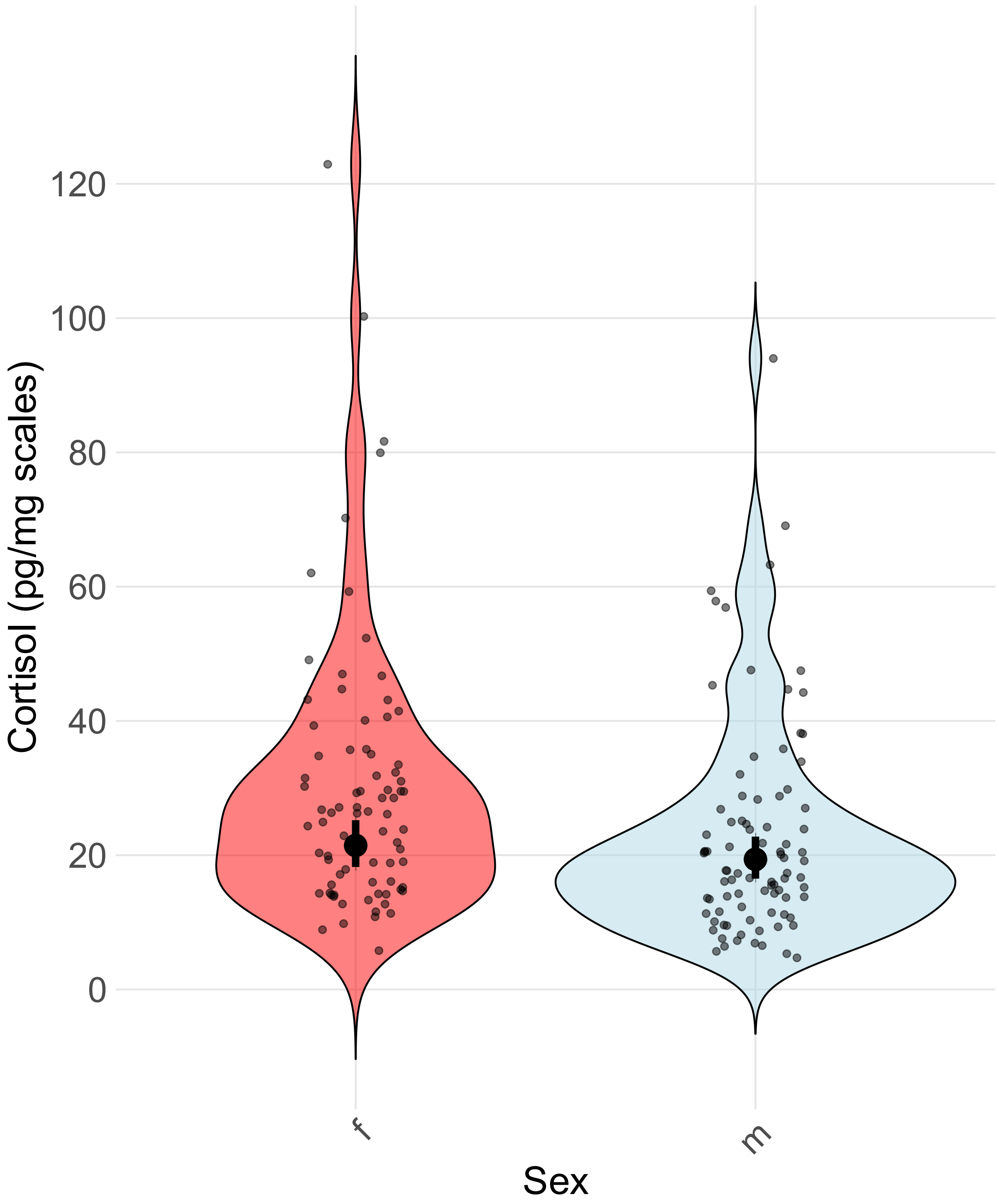

**Supplemental Figure 8.** Cortisol levels by sex based on model 2 testing for population × year differences. Large points with error bars represent the estimated marginal mean from the model for females (f) and males (m). Light points and violins show the distribution of the raw data.

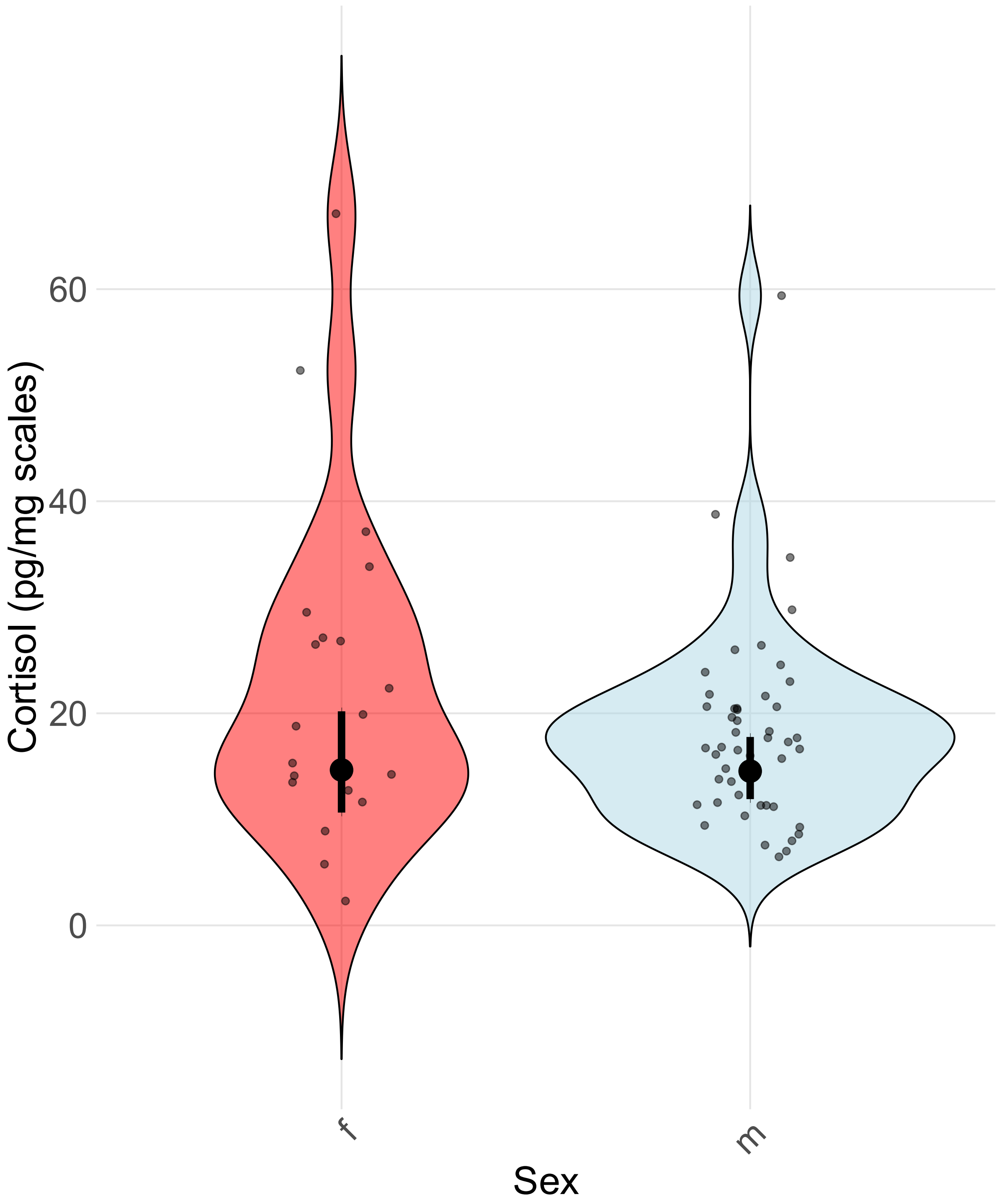

**Supplemental Figure 9.** Cortisol levels by sex based on model 3 testing for population × origin differences. Large points with error bars represent the estimated marginal mean from the model for females (f) and males (m). Light points and violins show the distribution of the raw data.

### Tables

**Supplemental Table 1.** Information about sampled individuals

| fish ID | population_year | origin | habitat | date_capture | date_dissection | time_dissection | date_hatched | sex | mass | standard_length | k_index |
|---|---|---|---|---|---|---|---|---|---|---|---|
| 1 | Pachon_23 | wild | cave | 05.04.23 | 05.04.23 | 2:00 PM | NA | m | 1.4190 | 4.1121 | 2.04[illegible] |
| 2 | Pachon_23 | wild | cave | 05.04.23 | 05.04.23 | 3:00 PM | NA | f | 3.2020 | 4.8346 | 2.83[illegible] |
| 3 | Pachon_23 | wild | cave | 05.04.23 | 05.04.23 | 4:00 PM | NA | m | 0.833 | 3.3518 | 2.21[illegible] |
| 4 | Pachon_23 | wild | cave | 05.04.23 | 05.04.23 | 4:40 PM | NA | f | 0.64 | 2.9471 | 2.50[illegible] |
| 5 | Pachon_23 | wild | cave | 05.04.23 | 05.04.23 | 5:30 PM | NA | m | 1.2330 | 3.8974 | 2.08[illegible] |
| 6 | Pachon_23 | wild | cave | 05.04.23 | 05.04.23 | 6:15 PM | NA | m | 1.5750 | 4.2972 | 1.98[illegible] |
| 7 | Pachon_23 | wild | cave | 05.04.23 | 05.04.23 | 6:30 PM | NA | m | 0.725 | 3.1976 | 2.21[illegible] |
| 8 | Pachon_23 | wild | cave | 05.04.23 | 05.04.23 | 7:20 PM | NA | NA | 0.725 | 3.2494 | 2.11[illegible] |
| 9 | Pachon_23 | wild | cave | 05.04.23 | 05.04.23 | 7:45 PM | NA | f | 1.1080 | 3.5795 | 2.41[illegible] |
| 10 | Pachon_23 | wild | cave | 05.04.23 | 05.04.23 | 8:35 PM | NA | f | 1.4760 | 4.0617 | 2.20[illegible] |
| 11 | Pachon_23 | wild | cave | 06.04.23 | 06.04.23 | 4:00 PM | NA | f | 1.5150 | 4.3268 | 1.87[illegible] |
| 12 | Pachon_23 | wild | cave | 06.04.23 | 06.04.23 | 4:40 PM | NA | m | 1.3890 | 4.2816 | 1.77[illegible] |
| 13 | Pachon_23 | wild | cave | 06.04.23 | 06.04.23 | 5:20 PM | NA | f | 1.1640 | 3.6472 | 2.39[illegible] |
| 14 | Pachon_23 | wild | cave | 06.04.23 | 06.04.23 | 6:00 PM | NA | m | 0.918 | 3.5133 | 2.11[illegible] |
| 15 | Pachon_23 | wild | cave | 06.04.23 | 06.04.23 | 6:45 PM | NA | m | 0.966 | 3.5346 | 2.18[illegible] |
| 16 | Pachon_23 | wild | cave | 06.04.23 | 06.04.23 | 7:00 PM | NA | m | 1.0210 | 3.5632 | 2.25[illegible] |
| 17 | Pachon_23 | wild | cave | 06.04.23 | 06.04.23 | 7:35 PM | NA | f | 1.2700 | 4.0965 | 1.84[illegible] |
| 18 | Pachon_23 | wild | cave | 06.04.23 | 06.04.23 | 8:15 PM | NA | m | 0.794 | 3.4804 | 1.88[illegible] |
| 19 | Pachon_23 | wild | cave | 06.04.23 | 06.04.23 | 8:50 PM | NA | m | 1.0230 | 3.6967 | 2.02[illegible] |
| 20 | Pachon_23 | wild | cave | 06.04.23 | 06.04.23 | 9:30 PM | NA | f | 0.779 | 3.4294 | 1.93[illegible] |
| 26 | Rio_Subterraneo_23 | wild | cave | 07.04.23 | 07.04.23 | 6:00 PM | NA | m | 3.7700 | 5.5766 | 2.17[illegible] |
| 27 | Rio_Subterraneo_23 | wild | cave | 07.04.23 | 07.04.23 | 7:05 PM | NA | f | 5.7700 | 6.0266 | 2.63[illegible] |
| 28 | Rio_Subterraneo_23 | wild | cave | 07.04.23 | 07.04.23 | 8:00 PM | NA | f | 5.5620 | 6.1829 | 2.35[illegible] |
| 29 | Rio_Subterraneo_23 | wild | cave | 07.04.23 | 07.04.23 | 9:00 PM | NA | m | 3.3090 | 5.1948 | 2.36[illegible] |
| 30 | Rio_Subterraneo_23 | wild | cave | 07.04.23 | 07.04.23 | 9:30 PM | NA | m | 1.1510 | 3.8356 | 2.04[illegible] |
| 31 | Rio_Subterraneo_23 | wild | cave | 07.04.23 | 07.04.23 | 10:05 PM | NA | f | 3.5020 | 5.1181 | 2.61[illegible] |
| 32 | Rio_Subterraneo_23 | wild | cave | 07.04.23 | 07.04.23 | 10:50 PM | NA | m | 1.0920 | 3.6291 | 2.28[illegible] |
| 33 | Rio_Subterraneo_23 | wild | cave | 07.04.23 | 07.04.23 | 11:15 PM | NA | m | 2.1280 | 4.3965 | 2.50[illegible] |
| 34 | Rio_Subterraneo_23 | wild | cave | 07.04.23 | 07.04.23 | 11:45 PM | NA | f | 1.2350 | 3.7453 | 2.35[illegible] |
| 35 | Rio_Subterraneo_23 | wild | cave | 07.04.23 | 08.04.23 | 12:30 AM | NA | m | 0.915 | 3.4134 | 2.30[illegible] |
| 36 | Rio_Subterraneo_23 | wild | cave | 08.04.23 | 08.04.23 | 2:10 PM | NA | m | 1.3010 | 3.7382 | 2.49[illegible] |
| 37 | Rio_Subterraneo_23 | wild | cave | 08.04.23 | 08.04.23 | 2:50 PM | NA | m | 1.6210 | 3.9956 | 2.54[illegible] |
| 38 | Rio_Subterraneo_23 | wild | cave | 08.04.23 | 08.04.23 | 3:30 PM | NA | f | 3.1400 | 5.4603 | 1.92[illegible] |
| 39 | Rio_Subterraneo_23 | wild | cave | 08.04.23 | 08.04.23 | 4:10 PM | NA | m | 1.9040 | 4.6609 | 1.88[illegible] |
| 40 | Rio_Subterraneo_23 | wild | cave | 08.04.23 | 08.04.23 | 4:45 PM | NA | f | 1.5890 | 4.4127 | 1.84[illegible] |
| 41 | Rio_Subterraneo_23 | wild | cave | 08.04.23 | 08.04.23 | 5:20 PM | NA | m | 1.2890 | 3.896 | 2.18[illegible] |
| 42 | Rio_Subterraneo_23 | wild | cave | 08.04.23 | 08.04.23 | 6:00 PM | NA | m | 1.7010 | 4.1793 | 2.33[illegible] |
| 43 | Rio_Subterraneo_23 | wild | cave | 08.04.23 | 08.04.23 | 6:45 PM | NA | f | 3.9070 | 5.3603 | 2.53[illegible] |

| | | | | | | | | | | | |
|---|---|---|---|---|---|---|---|---|---|---|---|
| **44** | Rio_Subterraneo_23 | wild | cave | 08.04.23 | 08.04.23 | 7:25 PM | NA | f | 3.9020 | 5.4603 | 2.397 |
| **45** | Rio_Subterraneo_23 | wild | cave | 08.04.23 | 08.04.23 | 8:05 PM | NA | m | 1.4010 | 3.8287 | 2.490 |
| **46** | Rio_Subterraneo_23 | wild | cave | 08.04.23 | 08.04.23 | 8:45 PM | NA | m | 1.2690 | 3.7124 | 2.480 |
| **47** | Rio_Subterraneo_23 | wild | cave | 08.04.23 | 08.04.23 | 9:20 PM | NA | m | 1.7460 | 4.6627 | 1.722 |
| **48** | Rio_Subterraneo_23 | wild | cave | 08.04.23 | 08.04.23 | 9:55 PM | NA | m | 0.772 | 3.48 | 1.832 |
| **49** | Rio_Subterraneo_23 | wild | cave | 08.04.23 | 08.04.23 | 10:30 PM | NA | m | 2.1980 | 5.1942 | 1.568 |
| **50** | Rio_Subterraneo_23 | wild | cave | 08.04.23 | 08.04.23 | 11:05 PM | NA | m | 1.0350 | 4.0619 | 1.544 |
| **51** | Rio_Choy_23 | wild | surface | 09.04.23 | 09.04.23 | 12:35 PM | NA | m | 6.8630 | 7.1601 | 1.870 |
| **52** | Rio_Choy_23 | wild | surface | 09.04.23 | 09.04.23 | 1:15 PM | NA | m | 9.1780 | 8.0899 | 1.733 |
| **53** | Rio_Choy_23 | wild | surface | 09.04.23 | 09.04.23 | 2:00 PM | NA | m | 6.5990 | 6.9257 | 1.980 |
| **54** | Rio_Choy_23 | wild | surface | 09.04.23 | 09.04.23 | 2:45 PM | NA | f | 4.7470 | 6.53 | 1.705 |
| **55** | Rio_Choy_23 | wild | surface | 09.04.23 | 09.04.23 | 3:30 PM | NA | f | 3.4340 | 5.4177 | 2.160 |
| **56** | Rio_Choy_23 | wild | surface | 09.04.23 | 09.04.23 | 3:50 PM | NA | m | 6.1600 | 6.5751 | 2.167 |
| **57** | Rio_Choy_23 | wild | surface | 09.04.23 | 09.04.23 | 4:20 PM | NA | m | 2.2230 | 4.845 | 1.955 |
| **58** | Rio_Choy_23 | wild | surface | 09.04.23 | 09.04.23 | 4:50 PM | NA | m | 4.9950 | 6.5328 | 1.792 |
| **59** | Rio_Choy_23 | wild | surface | 09.04.23 | 09.04.23 | 5:25 PM | NA | m | 1.3910 | 4.083 | 2.044 |
| **60** | Rio_Choy_23 | wild | surface | 09.04.23 | 09.04.23 | 6:00 PM | NA | f | 2.9040 | 5.198 | 2.068 |
| **61** | Rio_Choy_23 | wild | surface | 09.04.23 | 09.04.23 | 6:35 PM | NA | m | 1.1180 | 3.8295 | 1.991 |
| **62** | Rio_Choy_23 | wild | surface | 09.04.23 | 09.04.23 | 7:05 PM | NA | m | 5.8050 | 6.5772 | 2.040 |
| **63** | Rio_Choy_23 | wild | surface | 10.04.23 | 10.04.23 | 12:30 PM | NA | m | 5.8580 | 6.7445 | 1.909 |
| **64** | Rio_Choy_23 | wild | surface | 10.04.23 | 10.04.23 | 1:10 PM | NA | m | 7.5980 | 7.3712 | 1.897 |
| **65** | Rio_Choy_23 | wild | surface | 10.04.23 | 10.04.23 | 2:00 PM | NA | m | 5.1530 | 6.616 | 1.779 |
| **66** | Rio_Choy_23 | wild | surface | 10.04.23 | 10.04.23 | 2:35 PM | NA | m | 1.6060 | 4.5277 | 1.730 |
| **67** | Rio_Choy_23 | wild | surface | 10.04.23 | 10.04.23 | 3:10 PM | NA | m | 0.997 | 3.7788 | 1.848 |
| **68** | Rio_Choy_23 | wild | surface | 10.04.23 | 10.04.23 | 3:40 PM | NA | m | 1.2100 | 3.9321 | 1.990 |
| **69** | Rio_Choy_23 | wild | surface | 10.04.23 | 10.04.23 | 4:15 PM | NA | m | 1.3900 | 4.264 | 1.793 |
| **70** | Rio_Choy_23 | wild | surface | 10.04.23 | 10.04.23 | 4:45 PM | NA | m | 1.2900 | 4.2281 | 1.707 |
| **71** | Rio_Choy_23 | wild | surface | 10.04.23 | 10.04.23 | 5:10 PM | NA | m | 4.6640 | 6.1756 | 1.980 |
| **72** | Rio_Choy_23 | wild | surface | 10.04.23 | 10.04.23 | 5:40 PM | NA | m | 6.9520 | 7.2003 | 1.862 |
| **73** | Rio_Choy_23 | wild | surface | 10.04.23 | 10.04.23 | 6:20 PM | NA | m | 8.2440 | 7.7579 | 1.760 |
| **74** | Rio_Choy_23 | wild | surface | 10.04.23 | 10.04.23 | 6:50 PM | NA | m | 5.0270 | 6.5252 | 1.809 |
| **75** | Rio_Choy_23 | wild | surface | 10.04.23 | 10.04.23 | 7:25 PM | NA | f | 2.0690 | 4.6659 | 2.037 |
| **76** | Los_Sabinos_23 | wild | cave | 12.04.23 | 12.04.23 | 4:30 PM | NA | NA | 0.843 | 3.679 | 1.609 |
| **77** | Los_Sabinos_23 | wild | cave | 12.04.23 | 12.04.23 | 5:05 PM | NA | m | 1.6050 | 4.3971 | 1.888 |
| **78** | Los_Sabinos_23 | wild | cave | 12.04.23 | 12.04.23 | 5:50 PM | NA | m | 1.6540 | 4.3799 | 1.969 |
| **79** | Los_Sabinos_23 | wild | cave | 12.04.23 | 12.04.23 | 6:40 PM | NA | m | 1.8850 | 4.5288 | 2.029 |
| **80** | Los_Sabinos_23 | wild | cave | 12.04.23 | 12.04.23 | 7:15 PM | NA | m | 1.6310 | 4.2297 | 2.155 |
| **81** | Los_Sabinos_23 | wild | cave | 12.04.23 | 12.04.23 | 7:55 PM | NA | m | 2.9420 | 5.1287 | 2.181 |
| **82** | Los_Sabinos_23 | wild | cave | 12.04.23 | 12.04.23 | 8:45 PM | NA | m | 1.9370 | 4.761 | 1.795 |
| **83** | Los_Sabinos_23 | wild | cave | 12.04.23 | 12.04.23 | 9:30 PM | NA | m | 1.5150 | 4.0325 | 2.310 |
| **84** | Los_Sabinos_23 | wild | cave | 12.04.23 | 12.04.23 | 10:05 PM | NA | m | 1.1740 | 3.9127 | 1.960 |
| **85** | Los_Sabinos_23 | wild | cave | 12.04.23 | 12.04.23 | 10:40 PM | NA | m | 1.0900 | 4.146 | 1.529 |

| | | | | | | | | | | | |
|---|---|---|---|---|---|---|---|---|---|---|---|
| **86** | Los_Sabinos_23 | wild | cave | 12.04.23 | 12.04.23 | 11:20 PM | NA | m | 2.2100 | 5.0673 | 1.69[illegible] |
| **87** | Los_Sabinos_23 | wild | cave | 12.04.23 | 13.04.23 | 12:00 AM | NA | NA | 0.831 | 3.4804 | 1.97[illegible] |
| **88** | Los_Sabinos_23 | wild | cave | 12.04.23 | 13.04.23 | 9:30 AM | NA | m | 2.5250 | 5.4944 | 1.52[illegible] |
| **89** | Los_Sabinos_23 | wild | cave | 12.04.23 | 13.04.23 | 10:15 AM | NA | NA | 0.644 | 3.6621 | 1.31[illegible] |
| **90** | Los_Sabinos_23 | wild | cave | 12.04.23 | 13.04.23 | 10:55 AM | NA | NA | 0.88 | 3.6659 | 1.78[illegible] |
| **91** | Los_Sabinos_23 | wild | cave | 12.04.23 | 13.04.23 | 11:35 AM | NA | m | 2.1920 | 5.1272 | 1.62[illegible] |
| **92** | Los_Sabinos_23 | wild | cave | 12.04.23 | 13.04.23 | 12:20 PM | NA | f | 0.92 | 3.713 | 1.79[illegible] |
| **93** | Los_Sabinos_23 | wild | cave | 12.04.23 | 13.04.23 | 1:00 PM | NA | m | 2.8010 | 5.1142 | 2.09[illegible] |
| **94** | Los_Sabinos_23 | wild | cave | 12.04.23 | 13.04.23 | 1:50 PM | NA | m | 1.0020 | 4.0341 | 1.52[illegible] |
| **95** | Los_Sabinos_23 | wild | cave | 12.04.23 | 13.04.23 | 2:30 PM | NA | m | 1.3860 | 4.7137 | 1.32[illegible] |
| **96** | Los_Sabinos_23 | wild | cave | 12.04.23 | 13.04.23 | 3:00 PM | NA | m | 2.7560 | 5.1773 | 1.98[illegible] |
| **97** | Los_Sabinos_23 | wild | cave | 12.04.23 | 13.04.23 | 3:30 PM | NA | f | 1.6220 | 4.7457 | 1.51[illegible] |
| **98** | Los_Sabinos_23 | wild | cave | 12.04.23 | 13.04.23 | 4:10 PM | NA | m | 1.0420 | 4.03 | 1.59[illegible] |
| **99** | Los_Sabinos_23 | wild | cave | 12.04.23 | 13.04.23 | 4:40 PM | NA | m | 2.1540 | 4.8458 | 1.89[illegible] |
| **100** | Los_Sabinos_23 | wild | cave | 12.04.23 | 13.04.23 | 5:20 PM | NA | f | 2.3390 | 4.894 | 1.99[illegible] |
| **101** | Presa_El_Oyul_23 | wild | surface | 14.04.23 | 14.04.23 | 1:10 PM | NA | m | 1.8280 | 4.2629 | 2.36[illegible] |
| **102** | Presa_El_Oyul_23 | wild | surface | 14.04.23 | 14.04.23 | 1:50 PM | NA | m | 1.6220 | 4.4412 | 1.85[illegible] |
| **103** | Presa_El_Oyul_23 | wild | surface | 14.04.23 | 14.04.23 | 2:25 PM | NA | f | 2.7110 | 5.0964 | 2.04[illegible] |
| **104** | Presa_El_Oyul_23 | wild | surface | 14.04.23 | 14.04.23 | 3:00 PM | NA | f | 4.5350 | 6.0595 | 2.03[illegible] |
| **105** | Presa_El_Oyul_23 | wild | surface | 14.04.23 | 14.04.23 | 3:45 PM | NA | f | 2.3600 | 5.1778 | 1.70[illegible] |
| **106** | Presa_El_Oyul_23 | wild | surface | 14.04.23 | 14.04.23 | 4:45 PM | NA | f | 4.9110 | 6.1794 | 2.08[illegible] |
| **107** | Presa_El_Oyul_23 | wild | surface | 14.04.23 | 14.04.23 | 5:15 PM | NA | m | 1.5710 | 4.4631 | 1.76[illegible] |
| **108** | Presa_El_Oyul_23 | wild | surface | 14.04.23 | 14.04.23 | 5:45 PM | NA | f | 2.0260 | 4.711 | 1.93[illegible] |
| **109** | Presa_El_Oyul_23 | wild | surface | 14.04.23 | 14.04.23 | 6:25 PM | NA | f | 2.1150 | 4.8756 | 1.82[illegible] |
| **110** | Presa_El_Oyul_23 | wild | surface | 14.04.23 | 14.04.23 | 7:10 PM | NA | m | 2.5700 | 5.1612 | 1.86[illegible] |
| **111** | Presa_El_Oyul_23 | wild | surface | 14.04.23 | 14.04.23 | 8:10 PM | NA | m | 2.5430 | 5.1769 | 1.83[illegible] |
| **112** | Presa_El_Oyul_23 | wild | surface | 14.04.23 | 14.04.23 | 8:45 PM | NA | m | 1.4450 | 4.3573 | 1.74[illegible] |
| **113** | Presa_El_Oyul_23 | wild | surface | 14.04.23 | 14.04.23 | 9:10 PM | NA | f | 1.5410 | 4.2792 | 1.96[illegible] |
| **114** | Presa_El_Oyul_23 | wild | surface | 14.04.23 | 14.04.23 | 9:50 PM | NA | f | 2.8870 | 5.4433 | 1.79[illegible] |
| **115** | Presa_El_Oyul_23 | wild | surface | 15.04.23 | 15.04.23 | 11:50 AM | NA | m | 1.5540 | 4.213 | 2.07[illegible] |
| **116** | Presa_El_Oyul_23 | wild | surface | 15.04.23 | 15.04.23 | 12:25 PM | NA | f | 1.2960 | 3.8964 | 2.19[illegible] |
| **117** | Presa_El_Oyul_23 | wild | surface | 15.04.23 | 15.04.23 | 12:50 PM | NA | f | 2.3980 | 4.9707 | 1.95[illegible] |
| **118** | Presa_El_Oyul_23 | wild | surface | 15.04.23 | 15.04.23 | 1:25 PM | NA | f | 1.5690 | 4.2993 | 1.97[illegible] |
| **119** | Presa_El_Oyul_23 | wild | surface | 15.04.23 | 15.04.23 | 2:00 PM | NA | f | 2.1250 | 4.7183 | 2.02[illegible] |
| **120** | Presa_El_Oyul_23 | wild | surface | 15.04.23 | 15.04.23 | 2:30 PM | NA | f | 3.1800 | 5.3318 | 2.09[illegible] |
| **121** | Presa_El_Oyul_23 | wild | surface | 15.04.23 | 15.04.23 | 3:00 PM | NA | m | 2.1380 | 4.8107 | 1.92[illegible] |
| **122** | Presa_El_Oyul_23 | wild | surface | 15.04.23 | 15.04.23 | 3:30 PM | NA | f | 4.3810 | 6.2599 | 1.78[illegible] |
| **123** | Presa_El_Oyul_23 | wild | surface | 15.04.23 | 15.04.23 | 4:00 PM | NA | m | 2.7340 | 5.2768 | 1.86[illegible] |
| **124** | Presa_El_Oyul_23 | wild | surface | 15.04.23 | 15.04.23 | 4:30 PM | NA | f | 1.5000 | 4.2781 | 1.91[illegible] |
| **125** | Presa_El_Oyul_23 | wild | surface | 15.04.23 | 15.04.23 | 5:00 PM | NA | f | 4.3820 | 6.0682 | 1.96[illegible] |
| **126** | Presa_El_Oyul_24 | wild | surface | 09.03.24 | 09.03.24 | 12:55 PM | NA | f | 3.6050 | 5.658 | 1.99[illegible] |
| **127** | Presa_El_Oyul_24 | wild | surface | 09.03.24 | 09.03.24 | 1:42 PM | NA | f | 2.1840 | 4.543 | 2.32[illegible] |

| | | | | | | | | | | | |
|---|---|---|---|---|---|---|---|---|---|---|---|
| **128** | Presa_El_Oyul_24 | wild | surface | 09.03.24 | 09.03.24 | 2:21 PM | NA | f | 1.7740 | 4.417 | 2.059 |
| **129** | Presa_El_Oyul_24 | wild | surface | 09.03.24 | 09.03.24 | 2:51 PM | NA | f | 2.7290 | 5.019 | 2.158 |
| **130** | Presa_El_Oyul_24 | wild | surface | 09.03.24 | 09.03.24 | 3:23 PM | NA | f | 3.7620 | 5.603 | 2.139 |
| **131** | Presa_El_Oyul_24 | wild | surface | 09.03.24 | 09.03.24 | 3:56 PM | NA | f | 2.9070 | 5.110 | 2.179 |
| **132** | Presa_El_Oyul_24 | wild | surface | 09.03.24 | 09.03.24 | 4:30 PM | NA | f | 1.7780 | 4.400 | 2.087 |
| **133** | Presa_El_Oyul_24 | wild | surface | 09.03.24 | 09.03.24 | 4:57 PM | NA | f | 4.5000 | 5.997 | 2.080 |
| **134** | Presa_El_Oyul_24 | wild | surface | 09.03.24 | 09.03.24 | 5:31 PM | NA | f | 2.9460 | 5.201 | 2.094 |
| **135** | Presa_El_Oyul_24 | wild | surface | 09.03.24 | 09.03.24 | 6:00 PM | NA | f | 3.1860 | 5.406 | 2.017 |
| **136** | Presa_El_Oyul_24 | wild | surface | 09.03.24 | 09.03.24 | 6:28 PM | NA | f | 3.6420 | 5.548 | 2.133 |
| **137** | Presa_El_Oyul_24 | wild | surface | 09.03.24 | 09.03.24 | 6:56 PM | NA | f | 2.7640 | 5.209 | 1.950 |
| **138** | Presa_El_Oyul_24 | wild | surface | 09.03.24 | 09.03.24 | 7:30 PM | NA | f | 1.8890 | 4.553 | 2.001 |
| **139** | Presa_El_Oyul_24 | wild | surface | 09.03.24 | 10.03.24 | 10:57 AM | NA | f | 3.0000 | 5.267 | 2.053 |
| **140** | Presa_El_Oyul_24 | wild | surface | 09.03.24 | 10.03.24 | 11:20 AM | NA | f | 3.8520 | 5.643 | 2.144 |
| **141** | Presa_El_Oyul_24 | wild | surface | 09.03.24 | 10.03.24 | 11:45 AM | NA | f | 1.4520 | 4.118 | 2.079 |
| **142** | Presa_El_Oyul_24 | wild | surface | 09.03.24 | 10.03.24 | 12:10 PM | NA | f | 2.7230 | 5.113 | 2.037 |
| **143** | Presa_El_Oyul_24 | wild | surface | 09.03.24 | 10.03.24 | 12:27 PM | NA | f | 3.9370 | 5.846 | 1.971 |
| **144** | Presa_El_Oyul_24 | wild | surface | 09.03.24 | 10.03.24 | 12:49 PM | NA | f | 2.7720 | 5.100 | 2.090 |
| **145** | Presa_El_Oyul_24 | wild | surface | 09.03.24 | 10.03.24 | 1:15 PM | NA | f | 4.2330 | 5.926 | 2.034 |
| **146** | Presa_El_Oyul_24 | wild | surface | 09.03.24 | 10.03.24 | 1:44 PM | NA | f | 5.4340 | 6.402 | 2.071 |
| **147** | Presa_El_Oyul_24 | wild | surface | 09.03.24 | 10.03.24 | 2:12 PM | NA | f | 3.7060 | 5.605 | 2.105 |
| **148** | Presa_El_Oyul_24 | wild | surface | 09.03.24 | 10.03.24 | 2:36 PM | NA | f | 2.6020 | 5.011 | 2.068 |
| **149** | Presa_El_Oyul_24 | wild | surface | 09.03.24 | 10.03.24 | 3:00 PM | NA | f | 2.4160 | 5.001 | 1.932 |
| **150** | Presa_El_Oyul_24 | wild | surface | 09.03.24 | 10.03.24 | 3:18 PM | NA | f | 3.9160 | 6 | 1.999 |
| **151** | Rio_Choy_24 | wild | surface | 11.03.24 | 11.03.24 | 1:21 PM | NA | f | 16.2490 | 9.784 | 1.735 |
| **152** | Rio_Choy_24 | wild | surface | 11.03.24 | 11.03.24 | 1:48 PM | NA | m | 3.6500 | 5.834 | 1.838 |
| **153** | Rio_Choy_24 | wild | surface | 11.03.24 | 11.03.24 | 2:18 PM | NA | f | 1.5870 | 4.250 | 2.067 |
| **154** | Rio_Choy_24 | wild | surface | 11.03.24 | 11.03.24 | 2:39 PM | NA | f | 13.4270 | 8.401 | 2.265 |
| **155** | Rio_Choy_24 | wild | surface | 11.03.24 | 11.03.24 | 3:06 PM | NA | m | 3.0850 | 5.652 | 1.709 |
| **156** | Rio_Choy_24 | wild | surface | 11.03.24 | 11.03.24 | 3:30 PM | NA | f | 4.1370 | 5.914 | 2.000 |
| **157** | Rio_Choy_24 | wild | surface | 11.03.24 | 11.03.24 | 4:00 PM | NA | m | 8.2750 | 7.778 | 1.759 |
| **158** | Rio_Choy_24 | wild | surface | 11.03.24 | 11.03.24 | 4:24 PM | NA | f | 3.3310 | 5.550 | 1.948 |
| **159** | Rio_Choy_24 | wild | surface | 11.03.24 | 11.03.24 | 4:48 PM | NA | f | 3.9390 | 5.734 | 2.089 |
| **160** | Rio_Choy_24 | wild | surface | 11.03.24 | 11.03.24 | 5:09 PM | NA | m | 4.9550 | 6.340 | 1.944 |
| **161** | Rio_Choy_24 | wild | surface | 11.03.24 | 11.03.24 | 5:26 PM | NA | m | 5.7790 | 6.734 | 1.892 |
| **162** | Rio_Choy_24 | wild | surface | 11.03.24 | 11.03.24 | 5:47 PM | NA | m | 1.8980 | 4.570 | 1.989 |
| **163** | Rio_Choy_24 | wild | surface | 11.03.24 | 12.03.24 | 10:23 AM | NA | f | 10.0140 | 7.782 | 2.125 |
| **164** | Rio_Choy_24 | wild | surface | 11.03.24 | 12.03.24 | 10:50 AM | NA | m | 3.7740 | 5.758 | 1.977 |
| **165** | Rio_Choy_24 | wild | surface | 11.03.24 | 12.03.24 | 11:12 AM | NA | m | 3.8940 | 5.940 | 1.858 |
| **166** | Rio_Choy_24 | wild | surface | 11.03.24 | 12.03.24 | 11:38 AM | NA | m | 4.7430 | 6.108 | 2.081 |
| **167** | Rio_Choy_24 | wild | surface | 11.03.24 | 12.03.24 | 11:55 AM | NA | m | 4.7300 | 6.259 | 1.929 |
| **168** | Rio_Choy_24 | wild | surface | 11.03.24 | 12.03.24 | 12:17 PM | NA | m | 6.7130 | 6.821 | 2.115 |
| **169** | Rio_Choy_24 | wild | surface | 11.03.24 | 12.03.24 | 12:41 PM | NA | m | 5.5460 | 6.637 | 1.897 |

| | | | | | | | | | | | | |
|---|---|---|---|---|---|---|---|---|---|---|---|---|
| **170** | Rio_Choy_24 | wild | surface | 11.03.24 | 12.03.24 | 1:07 PM | NA | | m | 4.6850 | 6.343 | 1.83[illegible] |
| **171** | Rio_Choy_24 | wild | surface | 11.03.24 | 12.03.24 | 1:25 PM | NA | | m | 4.8240 | 6.234 | 1.99[illegible] |
| **172** | Rio_Choy_24 | wild | surface | 11.03.24 | 12.03.24 | 1:48 PM | NA | | f | 3.7140 | 5.544 | 2.18[illegible] |
| **173** | Rio_Choy_24 | wild | surface | 11.03.24 | 12.03.24 | 2:08 PM | NA | | f | 7.3150 | 7.112 | 2.03[illegible] |
| **174** | Rio_Choy_24 | wild | surface | 11.03.24 | 12.03.24 | 2:50 PM | NA | | f | 3.9680 | 5.743 | 2.09[illegible] |
| **175** | Rio_Choy_24 | wild | surface | 11.03.24 | 12.03.24 | 3:10 PM | NA | | m | 3.4160 | 5.500 | 2.05[illegible] |
| **176** | Rio_Subterraneo_24 | wild | cave | 14.03.24 | 14.03.24 | 6:08 PM | NA | | f | 0.407 | 3.037 | 1.45[illegible] |
| **177** | Rio_Subterraneo_24 | wild | cave | 14.03.24 | 14.03.24 | 6:45 PM | NA | | m | 1.4080 | 4.121 | 2.01[illegible] |
| **178** | Rio_Subterraneo_24 | wild | cave | 14.03.24 | 14.03.24 | 7:20 PM | NA | | f | 2.1530 | 4.620 | 2.18[illegible] |
| **179** | Rio_Subterraneo_24 | wild | cave | 14.03.24 | 14.03.24 | 8:00 PM | NA | | m | 1.3950 | 4.033 | 2.12[illegible] |
| **180** | Rio_Subterraneo_24 | wild | cave | 14.03.24 | 14.03.24 | 8:29 PM | NA | | m | 1.0540 | 3.820 | 1.89[illegible] |
| **181** | Rio_Subterraneo_24 | wild | cave | 14.03.24 | 14.03.24 | 9:07 PM | NA | | m | 3.2130 | 4.934 | 2.67[illegible] |
| **182** | Rio_Subterraneo_24 | wild | cave | 14.03.24 | 14.03.24 | 9:47 PM | NA | | m | 2.7400 | 4.767 | 2.52[illegible] |
| **183** | Rio_Subterraneo_24 | wild | cave | 14.03.24 | 14.03.24 | 10:26 PM | NA | | f | 3.1520 | 4.679 | 3.07[illegible] |
| **184** | Rio_Subterraneo_24 | wild | cave | 14.03.24 | 14.03.24 | 10:59 PM | NA | | m | 2.3080 | 4.727 | 2.18[illegible] |
| **185** | Rio_Subterraneo_24 | wild | cave | 14.03.24 | 14.03.24 | 11:32 PM | NA | | m | 2.5830 | 4.991 | 2.07[illegible] |
| **186** | Rio_Subterraneo_24 | wild | cave | 14.03.24 | 15.03.24 | 11:53 AM | NA | | m | 1.4880 | 4.204 | 2.00[illegible] |
| **187** | Rio_Subterraneo_24 | wild | cave | 14.03.24 | 15.03.24 | 12:19 PM | NA | | m | 1.5680 | 4.087 | 2.29[illegible] |
| **188** | Rio_Subterraneo_24 | wild | cave | 14.03.24 | 15.03.24 | 12:52 PM | NA | | m | 1.0730 | 3.840 | 1.89[illegible] |
| **189** | Rio_Subterraneo_24 | wild | cave | 14.03.24 | 15.03.24 | 1:25 PM | NA | | m | 1.6240 | 4.587 | 1.68[illegible] |
| **190** | Rio_Subterraneo_24 | wild | cave | 14.03.24 | 15.03.24 | 1:52 PM | NA | | f | 0.311 | 2.906 | 1.26[illegible] |
| **191** | Rio_Subterraneo_24 | wild | cave | 14.03.24 | 15.03.24 | 3:48 PM | NA | | m | 3.6620 | 5.435 | 2.28[illegible] |
| **192** | Rio_Subterraneo_24 | wild | cave | 14.03.24 | 15.03.24 | 4:30 PM | NA | | m | 2.2600 | 4.800 | 2.04[illegible] |
| **193** | Rio_Subterraneo_24 | wild | cave | 14.03.24 | 15.03.24 | 4:57 PM | NA | | f | 4.0560 | 5.517 | 2.41[illegible] |
| **194** | Rio_Subterraneo_24 | wild | cave | 14.03.24 | 15.03.24 | 5:50 PM | NA | | m | 0.685 | 3.342 | 1.83[illegible] |
| **195** | Rio_Subterraneo_24 | wild | cave | 14.03.24 | 15.03.24 | 6:10 PM | NA | | f | 2.6160 | 4.678 | 2.55[illegible] |
| **201** | Tinaja_24 | wild | cave | 19.03.24 | 20.03.24 | 11:55 AM | NA | | f | 1.1410 | 4.031 | 1.74[illegible] |
| **202** | Tinaja_24 | wild | cave | 19.03.24 | 20.03.24 | 12:47 PM | NA | | f | 1.1230 | 3.958 | 1.81[illegible] |
| **203** | Tinaja_24 | wild | cave | 19.03.24 | 20.03.24 | 1:49 PM | NA | | f | 1.1330 | 3.795 | 2.07[illegible] |
| **204** | Tinaja_24 | wild | cave | 19.03.24 | 20.03.24 | 2:39 PM | NA | | f | 1.4620 | 4.39 | 1.72[illegible] |
| **205** | Tinaja_24 | wild | cave | 19.03.24 | 20.03.24 | 3:22 PM | NA | | f | 0.733 | 3.553 | 1.63[illegible] |
| **206** | Tinaja_24 | wild | cave | 19.03.24 | 20.03.24 | 4:15 PM | NA | | f | 2.6950 | 5.313 | 1.79[illegible] |
| **207** | Tinaja_24 | wild | cave | 19.03.24 | 20.03.24 | 4:51 PM | NA | | m | 1.5010 | 4.328 | 1.85[illegible] |
| **208** | Tinaja_24 | wild | cave | 19.03.24 | 20.03.24 | 6:00 PM | NA | | m | 1.5290 | 4.527 | 1.64[illegible] |
| **209** | Tinaja_24 | wild | cave | 19.03.24 | 20.03.24 | 6:42 PM | NA | | f | 1.0130 | 3.822 | 1.81[illegible] |
| **210** | Tinaja_24 | wild | cave | 19.03.24 | 20.03.24 | 7:28 PM | NA | | f | 1.5510 | 4.254 | 2.01[illegible] |
| **211** | Tinaja_24 | wild | cave | 19.03.24 | 20.03.24 | 8:02 PM | NA | | f | 0.807 | 3.671 | 1.63[illegible] |
| **221** | Pachon_lab | laboratory | cave | 08.05.24 | 08.05.24 | NA | | 22.02.23 | f | 2.0800 | 4.400 | 2.441773[illegible] |
| **222** | Pachon_lab | laboratory | cave | 08.05.24 | 08.05.24 | NA | | 22.02.23 | m | 2.0490 | 4.500 | 2.248559[illegible] |
| **223** | Pachon_lab | laboratory | cave | 08.05.24 | 08.05.24 | NA | | 22.02.23 | m | 1.9210 | 4.100 | 2.787249[illegible] |
| **224** | Pachon_lab | laboratory | cave | 08.05.24 | 08.05.24 | NA | | 22.02.23 | m | 1.7170 | 4.100 | 2.491258[illegible] |
| **225** | Pachon_lab | laboratory | cave | 08.05.24 | 08.05.24 | NA | | 22.02.23 | m | 1.7030 | 3.900 | 2.870918[illegible] |

| | | | | | | | | | | | |
|---|---|---|---|---|---|---|---|---|---|---|---|
| **226** | Pachon_lab | laboratory | cave | 08.05.24 | 08.05.24 | NA | 22.02.23 | m | 1.8270 | 4.300 | 2.2979109 |
| **227** | Pachon_lab | laboratory | cave | 08.05.24 | 08.05.24 | NA | 22.02.23 | m | 1.9480 | 4.300 | 2.4500987 |
| **228** | Pachon_lab | laboratory | cave | 08.05.24 | 08.05.24 | NA | 22.02.23 | m | 1.9580 | 4.200 | 2.6428032 |
| **229** | Pachon_lab | laboratory | cave | 14.05.24 | 14.05.24 | NA | 22.02.23 | NA | 1.7840 | 3.800 | 3.2512028 |
| **230** | Pachon_lab | laboratory | cave | 14.05.24 | 14.05.24 | NA | 22.02.23 | m | 1.8570 | 4.200 | 2.5064788 |
| **231** | Rio_Choy_lab | laboratory | surface | 14.05.24 | 14.05.24 | NA | 30.06.23 | f | 2.0310 | 4.400 | 2.3842500 |
| **232** | Rio_Choy_lab | laboratory | surface | 14.05.24 | 14.05.24 | NA | 30.06.23 | m | 0.843 | 3.400 | 2.1448199 |
| **233** | Rio_Choy_lab | laboratory | surface | 14.05.24 | 14.05.24 | NA | 30.06.23 | m | 0.98 | 4.200 | 1.3227513 |
| **234** | Rio_Choy_lab | laboratory | surface | 14.05.24 | 14.05.24 | NA | 30.06.23 | m | 0.883 | 3.500 | 2.0594752 |
| **235** | Rio_Choy_lab | laboratory | surface | 14.05.24 | 14.05.24 | NA | 30.06.23 | f | 0.901 | 3.200 | 2.7496338 |
| **236** | Rio_Choy_lab | laboratory | surface | 14.05.24 | 14.05.24 | NA | 30.06.23 | f | 1.1760 | 3.600 | 2.5205761 |
| **237** | Rio_Choy_lab | laboratory | surface | 15.05.24 | 15.05.24 | NA | 30.06.23 | m | 0.612 | 3.100 | 2.0543117 |
| **238** | Rio_Choy_lab | laboratory | surface | 15.05.24 | 15.05.24 | NA | 30.06.23 | f | 1.8500 | 4.200 | 2.4970300 |
| **239** | Rio_Choy_lab | laboratory | surface | 15.05.24 | 15.05.24 | NA | 30.06.23 | m | 1.2250 | 4.000 | 1.9140625 |
| **240** | Rio_Choy_lab | laboratory | surface | 15.05.24 | 15.05.24 | NA | 30.06.23 | f | 0.737 | 3.200 | 2.2491455 |
| **251** | Tinaja_23 | wild | cave | 02.12.23 | 02.12.23 | NA | NA | NA | 0.9 | 3.385 | 2.3204194 |
| **252** | Tinaja_23 | wild | cave | 02.12.23 | 02.12.23 | NA | NA | NA | 0.4 | 3.070 | 1.3824355 |
| **253** | Tinaja_23 | wild | cave | 02.12.23 | 02.12.23 | NA | NA | f | 2.8000 | 5.274 | 1.9086999 |
| **254** | Tinaja_23 | wild | cave | 02.12.23 | 02.12.23 | NA | NA | m | 1.5000 | 4.049 | 2.2596851 |
| **255** | Tinaja_23 | wild | cave | 02.12.23 | 02.12.23 | NA | NA | m | 2.1000 | 4.903 | 1.781690 |
| **256** | Tinaja_23 | wild | cave | 02.12.23 | 02.12.23 | NA | NA | m | 2.0000 | 4.899 | 1.7010132 |
| **257** | Tinaja_23 | wild | cave | 02.12.23 | 02.12.23 | NA | NA | m | 0.9 | 3.728 | 1.7370599 |
| **258** | Tinaja_23 | wild | cave | 02.12.23 | 02.12.23 | NA | NA | m | 1.5000 | 4.272 | 1.9239660 |
| **259** | Tinaja_23 | wild | cave | 02.12.23 | 02.12.23 | NA | NA | f | 3.6000 | 5.862 | 1.7871667 |
| **260** | Tinaja_23 | wild | cave | 02.12.23 | 02.12.23 | NA | NA | m | 2.8000 | 5.609 | 1.5867252 |
| **261** | Tinaja_23 | wild | cave | 02.12.23 | 02.12.23 | NA | NA | m | 2.4000 | 5.526 | 1.4222587 |
| **262** | Tinaja_23 | wild | cave | 02.12.23 | 02.12.23 | NA | NA | m | 2.5000 | 5.352 | 1.6307662 |
| **263** | Tinaja_23 | wild | cave | 02.12.23 | 02.12.23 | NA | NA | f | 3.6000 | 5.778 | 1.8662503 |
| **264** | Tinaja_23 | wild | cave | 02.12.23 | 02.12.23 | NA | NA | m | 2.0000 | 5.037 | 1.5649993 |
| **265** | Tinaja_23 | wild | cave | 03.12.23 | 03.12.23 | NA | NA | f | 1.1000 | 3.908 | 1.8430150 |
| **266** | Tinaja_23 | wild | cave | 03.12.23 | 03.12.23 | NA | NA | f | 1.0000 | 3.820 | 1.7939483 |
| **267** | Tinaja_23 | wild | cave | 03.12.23 | 03.12.23 | NA | NA | NA | 0.6 | 3.186 | 1.8552991 |
| **268** | Tinaja_23 | wild | cave | 03.12.23 | 03.12.23 | NA | NA | f | 1.4000 | 4.326 | 1.7292920 |
| **269** | Tinaja_23 | wild | cave | 03.12.23 | 03.12.23 | NA | NA | m | 0.7 | 3.674 | 1.411499 |
| **270** | Tinaja_23 | wild | cave | 03.12.23 | 03.12.23 | NA | NA | NA | 0.8 | 3.532 | 1.8156323 |
| **271** | Tinaja_23 | wild | cave | 03.12.23 | 03.12.23 | NA | NA | NA | 0.5 | 3.067 | 1.7331202 |
| **272** | Tinaja_23 | wild | cave | 03.12.23 | 03.12.23 | NA | NA | NA | 0.4 | 2.795 | 1.831954 |
| **273** | Tinaja_23 | wild | cave | 03.12.23 | 03.12.23 | NA | NA | NA | 0.6 | 3.137 | 1.9436030 |

**Supplemental Table 2.** Average water quality measurements over a period of one year in the housing tanks of Pachón cave and Rio Choy surface laboratory populations

| Parameter | Pachón tank | Rio Choy tank |
|---|---|---|
| pH | 8.523 | 8.550 |
| Cond ($\mu$S/cm) | 661.500 | 646.333 |
| NO2 (mg/L) | 0.007 | 0.008 |
| NH3 (mg/L) | 0.010 | 0.011 |
| PO4 (mg/L) | 0.883 | 0.400 |

**Supplemental Table 3.** Back-transformed (from log to normal) estimated marginal means (EMMs) of cortisol levels per habitat from model 1 with lower and upper confidence limits (CL) and standard error (SE).

| habitat | EMM | SE | lower.CL | upper.CL |
|---|---|---|---|---|
| cave | 21.451981 | 1.07564451 | 18.5754417 | 24.7739729 |
| lake | 37.1294475 | 1.10339122 | 30.5739371 | 45.0905577 |
| river | 21.5154886 | 1.10312275 | 17.7252659 | 26.1161808 |

**Supplemental Table 4.** Pairwise comparisons (contrast) of habitats in their scale cortisol levels based on model 1 with estimate, standard error (SE), degrees of freedom (df), t-statistic (t.ratio), adjusted p-value (p.value) and significance.

| contrast | estimate | SE | df | t.ratio | p.value | significance |
|---|---|---|---|---|---|---|
| cave - lake | -0.5485934 | 0.12732036 | 165 | -4.3087642 | 8.31E-05 | *** |
| cave - river | -0.0029561 | 0.12041795 | 165 | -0.0245485 | 0.99966781 | ns |
| lake - river | 0.54563732 | 0.14452544 | 165 | 3.77537211 | 0.00064893 | *** |

**Supplemental Table 5.** Back-transformed (from log to normal) estimated marginal means (EMMs) of cortisol levels per population and year from model 2 with lower and upper confidence limits (CL) and standard error (SE).

| population | year | EMM | SE | lower.CL | upper.CL |
|---|---|---|---|---|---|
| Los_Sabinos | 2023 | 13.0685184 | 1.13086013 | 10.2503542 | 16.66149 |
| Pachón | 2023 | 9.46815971 | 1.24463456 | 6.1453859 | 14.5875376 |
| Presa_El_Oyul | 2023 | 43.1916419 | 1.11679392 | 34.7255029 | 53.7218405 |
| Rio_Choy | 2023 | 20.9692267 | 1.126768 | 16.5654979 | 26.5436313 |
| Rio_Subterráneo | 2023 | 20.2139947 | 1.13125413 | 15.8440376 | 25.7892332 |
| Tinaja | 2023 | 20.9901964 | 1.24050122 | 13.7136614 | 32.1276959 |
| Presa_El_Oyul | 2024 | 29.3215224 | 1.12353465 | 23.2955826 | 36.9062105 |
| Rio_Choy | 2024 | 22.20855 | 1.1148895 | 17.9156677 | 27.530076 |

| | | | | | |
|---|---|---|---|---|---|
| Rio_Subterráneo | 2024 | 38.4573607 | 1.15221387 | 29.0700606 | 50.8760066 |
| Tinaja | 2024 | 24.852957 | 1.18878633 | 17.6620515 | 34.9715587 |

**Supplemental Table 6.** Pairwise comparisons (contrast) of population × year in their scale cortisol levels based on model 2 with estimate, standard error (SE), degrees of freedom (df), t-statistic (t.ratio), adjusted p-value (p.value) and significance.

| contrast | estimate | SE | df | t.ratio | p.value | significance |
|---|---|---|---|---|---|---|
| Los_Sabinos year2023 - Pachón year2023 | 0.32227 | 0.2561 | 158 | 1.25856 | 0.96112 | ns |
| Los_Sabinos year2023 - Presa_El_Oyul year2023 | -1.19544 | 0.166 | 158 | -7.20308 | 1.02E-09 | *** |
| Los_Sabinos year2023 - Rio_Choy year2023 | -0.47285 | 0.1621 | 158 | -2.91619 | 0.10979 | ns |
| Los_Sabinos year2023 - Rio_Subterráneo year2023 | -0.43617 | 0.1781 | 158 | -2.44853 | 0.30492 | ns |
| Los_Sabinos year2023 - Tinaja year2023 | -0.47385 | 0.2443 | 158 | -1.93925 | 0.64238 | ns |
| Los_Sabinos year2023 - Presa_El_Oyul year2024 | -0.80812 | 0.1788 | 158 | -4.52062 | 0.0005 | *** |
| Los_Sabinos year2023 - Rio_Choy year2024 | -0.53027 | 0.1612 | 158 | -3.29015 | 0.03956 | * |
| Los_Sabinos year2023 - Rio_Subterráneo year2024 | -1.07934 | 0.1895 | 158 | -5.69521 | 2.59E-06 | *** |
| Los_Sabinos year2023 - Tinaja year2024 | -0.64277 | 0.2118 | 158 | -3.03414 | 0.08103 | ns |
| Pachón year2023 - Presa_El_Oyul year2023 | -1.51771 | 0.249 | 158 | -6.09455 | 3.60E-07 | *** |
| Pachón year2023 - Rio_Choy year2023 | -0.79512 | 0.2527 | 158 | -3.14645 | 0.05973 | ns |
| Pachón year2023 - Rio_Subterráneo year2023 | -0.75844 | 0.2404 | 158 | -3.15505 | 0.05831 | ns |
| Pachón year2023 - Tinaja year2023 | -0.79612 | 0.3144 | 158 | -2.53191 | 0.25994 | ns |
| Pachón year2023 - Presa_El_Oyul year2024 | -1.13039 | 0.2482 | 158 | -4.55509 | 0.00043 | *** |
| Pachón year2023 - Rio_Choy year2024 | -0.85254 | 0.2456 | 158 | -3.4707 | 0.02281 | * |
| Pachón year2023 - Rio_Subterráneo year2024 | -1.40162 | 0.2494 | 158 | -5.62016 | 3.72E-06 | *** |
| Pachón year2023 - Tinaja year2024 | -0.96504 | 0.2881 | 158 | -3.34982 | 0.03311 | * |
| Presa_El_Oyul year2023 - Rio_Choy year2023 | 0.72259 | 0.1641 | 158 | 4.40219 | 0.00081 | *** |
| Presa_El_Oyul year2023 - Rio_Subterráneo year2023 | 0.75927 | 0.1715 | 158 | 4.42806 | 0.00073 | *** |
| Presa_El_Oyul year2023 - Tinaja year2023 | 0.72159 | 0.2393 | 158 | 3.01594 | 0.08502 | ns |
| Presa_El_Oyul year2023 - Presa_El_Oyul year2024 | 0.38733 | 0.1559 | 158 | 2.48523 | 0.28459 | ns |
| Presa_El_Oyul year2023 - Rio_Choy year2024 | 0.66517 | 0.1554 | 158 | 4.28021 | 0.0013 | ** |
| Presa_El_Oyul year2023 - Rio_Subterráneo year2024 | 0.1161 | 0.1865 | 158 | 0.62247 | 0.9998 | ns |
| Presa_El_Oyul year2023 - Tinaja year2024 | 0.55267 | 0.1991 | 158 | 2.77614 | 0.15384 | ns |
| Rio_Choy year2023 - Rio_Subterráneo year2023 | 0.03668 | 0.1736 | 158 | 0.21134 | 1 | ns |
| Rio_Choy year2023 - Tinaja year2023 | -0.001 | 0.2438 | 158 | -0.0041 | 1 | ns |
| Rio_Choy year2023 - Presa_El_Oyul year2024 | -0.33527 | 0.1765 | 158 | -1.89924 | 0.66957 | ns |

| | | | | | | |
|---|---|---|---|---|---|---|
| Rio_Choy year2023 - Rio_Choy year2024 | -0.05742 | 0.1586 | 158 | -0.36197 | 1 | ns |
| Rio_Choy year2023 - Rio_Subterráneo year2024 | -0.60649 | 0.185 | 158 | -3.27804 | 0.04099 | * |
| Rio_Choy year2023 - Tinaja year2024 | -0.16992 | 0.2115 | 158 | -0.80336 | 0.99845 | ns |
| Rio_Subterráneo year2023 - Tinaja year2023 | -0.03768 | 0.256 | 158 | -0.14718 | 1 | ns |
| Rio_Subterráneo year2023 - Presa_El_Oyul year2024 | -0.37195 | 0.1732 | 158 | -2.14705 | 0.49757 | ns |
| Rio_Subterráneo year2023 - Rio_Choy year2024 | -0.0941 | 0.1653 | 158 | -0.56929 | 0.99991 | ns |
| Rio_Subterráneo year2023 - Rio_Subterráneo year2024 | -0.64317 | 0.1726 | 158 | -3.72745 | 0.00983 | ** |
| Rio_Subterráneo year2023 - Tinaja year2024 | -0.2066 | 0.2242 | 158 | -0.92168 | 0.99556 | ns |
| Tinaja year2023 - Presa_El_Oyul year2024 | -0.33427 | 0.2448 | 158 | -1.3655 | 0.93579 | ns |
| Tinaja year2023 - Rio_Choy year2024 | -0.05642 | 0.2405 | 158 | -0.23464 | 1 | ns |
| Tinaja year2023 - Rio_Subterráneo year2024 | -0.60549 | 0.266 | 158 | -2.27658 | 0.4101 | ns |
| Tinaja year2023 - Tinaja year2024 | -0.16892 | 0.2694 | 158 | -0.62712 | 0.99979 | ns |
| Presa_El_Oyul year2024 - Rio_Choy year2024 | 0.27784 | 0.1625 | 158 | 1.70961 | 0.78851 | ns |
| Presa_El_Oyul year2024 - Rio_Subterráneo year2024 | -0.27123 | 0.1901 | 158 | -1.42701 | 0.91699 | ns |
| Presa_El_Oyul year2024 - Tinaja year2024 | 0.16535 | 0.2013 | 158 | 0.82133 | 0.99816 | ns |
| Rio_Choy year2024 - Rio_Subterráneo year2024 | -0.54907 | 0.1788 | 158 | -3.07113 | 0.07341 | ns |
| Rio_Choy year2024 - Tinaja year2024 | -0.1125 | 0.2045 | 158 | -0.55004 | 0.99993 | ns |
| Rio_Subterráneo year2024 - Tinaja year2024 | 0.43657 | 0.2371 | 158 | 1.84094 | 0.70815 | ns |

**Supplemental Table 7.** Back-transformed (from log to normal) estimated marginal means (EMMs) of cortisol levels per population and origin (laboratory vs. wild) from model 3 with lower and upper confidence limits (CL) and standard error (SE).

| population | origin | EMM | SE | lower.CL | upper.CL |
|---|---|---|---|---|---|
| Pachón | lab | 9.039 | 1.29868 | 5.360836 | 15.24069 |
| Rio_Choy | lab | 23.737 | 1.1816 | 17.00412 | 33.13539 |
| Pachón | wild | 9.5116 | 1.24922 | 6.096418 | 14.83981 |
| Rio_Choy | wild | 22.256 | 1.09282 | 18.63798 | 26.57746 |

**Supplemental Table 8.** Pairwise comparisons (contrast) of population × origin in their scale cortisol levels based on model 3 with estimate, standard error (SE), degrees of freedom (df), t-statistic (t.ratio), adjusted p-value (p.value) and significance.

| contrast | estimate | SE | df | t.ratio | p.value | significance |
|---|---|---|---|---|---|---|
| Pachón lab - Rio_Choy lab | -0.96548 | 0.287 | 62 | -3.3643 | 0.00705 | ** |
| Pachón lab - Pachón wild | -0.05096 | 0.3016 | 62 | -0.169 | 0.99827 | ns |
| Pachón lab - Rio_Choy wild | -0.90109 | 0.2891 | 62 | -3.1168 | 0.01435 | * |
| Rio_Choy lab - Pachón wild | 0.914522 | 0.2677 | 62 | 3.41592 | 0.00605 | ** |

| | | | | | | |
|---|---|---|---|---|---|---|
| Rio_Choy lab - Rio_Choy wild | 0.064397 | 0.1949 | 62 | 0.33039 | 0.98744 | ns |
| Pachón wild - Rio_Choy wild | -0.85013 | 0.2486 | 62 | -3.4196 | 0.00598 | ** |

**Supplemental Table 9.** Coefficients for model 1 with estimate, standard error, t value, p value (Pr(>|t|) and Signif. codes: 0 ‘***’ 0.001 ‘**’ 0.01 ‘*’ 0.05 ‘.’ 0.1 ‘ns’ 1.

| Coefficients | Estimate | Std. Error | t value | Pr(>\|t\|) | Significance |
|---|---|---|---|---|---|
| Intercept | 1.763833 | 0.417669 | 4.223 | 3.97E-05 | *** |
| habitatlake | 0.548593 | 0.12732 | 4.309 | 2.81E-05 | *** |
| habitatriver | 0.002956 | 0.120418 | 0.025 | 0.980445 | ns |
| k_index | 0.65162 | 0.190419 | 3.422 | 0.000784 | *** |
| sexm | -0.020383 | 0.109919 | -0.185 | 0.85311 | ns |

**Supplemental Table 10.** Coefficients for model 2 with estimate, standard error, t value, p value (Pr(>|t|) and Signif. codes: 0 ‘***’ 0.001 ‘**’ 0.01 ‘*’ 0.05 ‘.’ 0.1 ‘ns’ 1.

| Coefficients | Estimate | Std. Error | t value | Pr(>\|t\|) | Significa |
|---|---|---|---|---|---|
| Intercept | 1.6642 | 0.4075 | 4.084 | 7.02E-05 | *** |
| populationPachón | -0.3223 | 0.2561 | -1.259 | 0.21005 | ns |
| populationPresa_El_Oyul | 1.1954 | 0.166 | 7.203 | 2.26E-11 | *** |
| populationRio_Choy | 0.4728 | 0.1621 | 2.916 | 0.00406 | ** |
| populationRio_Subterráneo | 0.4362 | 0.1781 | 2.449 | 0.01544 | * |
| populationTinaja | 0.4738 | 0.2443 | 1.939 | 0.05425 | . |
| year2024 | 0.1689 | 0.2694 | 0.627 | 0.53149 | ns |
| k_index | 0.475 | 0.195 | 2.435 | 0.01599 | * |
| sexm | -0.1012 | 0.1002 | -1.009 | 0.31444 | ns |
| populationPachón:year2024 | NA | NA | NA | NA | NA |
| populationPresa_El_Oyul:year2024 | -0.5562 | 0.3083 | -1.804 | 0.07312 | . |
| populationRio_Choy:year2024 | -0.1115 | 0.3105 | -0.359 | 0.72002 | ns |
| populationRio_Subterráneo:year2024 | 0.4743 | 0.3211 | 1.477 | 0.14171 | ns |
| populationTinaja:year2024 | NA | NA | NA | NA | NA |

**Supplemental Table 11.** Coefficients for model 3 with estimate, standard error, t value, p value (Pr(>|t|) and Signif. codes: 0 ‘***’ 0.001 ‘**’ 0.01 ‘*’ 0.05 ‘.’ 0.1 ‘ns’ 1.

| Coefficients | Estimate | Std. Error | t value | Pr(>\|t\|) | Significance |
|---|---|---|---|---|---|
| Intercept | 0.891534 | 0.929126 | 0.96 | 0.34101 | ns |
| populationRio_Choy | 0.965485 | 0.286976 | 3.364 | 0.00132 | ** |
| originwild | 0.050963 | 0.301584 | 0.169 | 0.86636 | ns |
| k_index | 0.633455 | 0.33041 | 1.917 | 0.05983 | . |
| sexm | -0.007535 | 0.157927 | -0.048 | 0.9621 | ns |
| populationRio_Choy:originwild | -0.11536 | 0.343289 | -0.336 | 0.73797 | ns |

**Supplemental Table 12.** 95% confidence intervals based on bootstrap resampling with 1000 iterations per population and year/origin.

| Population_year/origin | Lower_CI | Upper_CI |
|---|---|---|

| | | |
|---|---|---|
| Pachon_23 | 3.15 | 59.78 |
| Rio_Subterraneo_23 | 55.96 | 282.39 |
| Rio_Choy_23 | 19.30 | 68.63 |
| Los_Sabinos_23 | 12.57 | 48.83 |
| Presa_El_Oyul_23 | 58.55 | 1018.53 |
| Presa_El_Oyul_24 | 104.87 | 954.44 |
| Rio_Choy_24 | 16.29 | 271.85 |
| Rio_Subterraneo_24 | 267.54 | 908.98 |
| Tinaja_24 | 21.12 | 75.14 |
| Pachon_lab | 10.25 | 93.99 |
| Rio_Choy_lab | 79.49 | 612.57 |
| Tinaja_23 | 15.55 | 94.95 |
